# Phosphorylation of CRYAB Induces a Condensatopathy to Worsen Post-Myocardial Infarction Left Ventricular Remodeling

**DOI:** 10.1101/2024.08.30.610556

**Authors:** Moydul Islam, David R. Rawnsley, Xiucui Ma, Walter Navid, Chen Zhao, Layla Foroughi, John T. Murphy, Honora Navid, Carla J. Weinheimer, Attila Kovacs, Jessica Nigro, Aaradhya Diwan, Ryan Chang, Minu Kumari, Martin E. Young, Babak Razani, Kenneth B. Margulies, Mahmoud Abdellatif, Simon Sedej, Ali Javaheri, Douglas F. Covey, Kartik Mani, Abhinav Diwan

**Affiliations:** Division of Cardiology and Center for Cardiovascular Research, Department of Medicine, Washington University School of Medicine, St. Louis, MO, USA; Center for Cardiovascular Research, Department of Medicine, Washington University School of Medicine, St. Louis, MO, USA; Department of Chemistry, Washington University in St. Louis, MO, USA; Division of Cardiology and Department of Medicine, University of Alabama, Birmingham, AL, USA; John Cochran Veterans Affairs Medical Center, St. Louis, MO, USA; Department of Medicine, University of Pittsburgh, Pittsburgh, PA, USA; Department of Medicine, University of Pennsylvania, Philadelphia, PA, USA; Division of Cardiology, Medical University of Graz, Graz Austria; BioTechMed-Graz, Graz, Austria; Institute of Physiology, University of Maribor, Maribor, Slovenia; Department of Developmental Biology, Washington University in St. Louis, MO, USA; Dept. of Anesthesiology, Psychiatry and Taylor Family Institute for Innovative Psychiatric Research, Washington University in St. Louis, MO, USA; Iowa City Veterans Affairs Medical Center and University of Iowa, Iowa City, IA; Departments of Cell Biology and Physiology, Obstetrics and Gynecology, and Neurology, Washington University in St. Louis, MO, USA

**Author notes:** Corresponding Author: Abhinav Diwan, MD, Professor of Medicine, Neurology, Cell Biology and Physiology, Obstetrics and Gynecology; Washington University School of Medicine, Division of Cardiology, 660 S. Euclid, CSRB 827 NTA, St. Louis, MO 63110; or Kartik Mani, MBBS, Clinical Associate Professor, University of Iowa and Iowa City VA Medical Center; 200 Hawkins Drive Iowa City, IA 52242. Contributed equally (Moydul Islam, David R. Rawnsley).

## Abstract

Protein aggregates are emerging therapeutic targets in rare monogenic causes of cardiomyopathy and amyloid heart disease, but their role in more prevalent heart failure syndromes remains mechanistically unexamined. We observed mis-localization of desmin and sarcomeric proteins to aggregates in human myocardium with ischemic cardiomyopathy and in mouse hearts with post-myocardial infarction ventricular remodeling, mimicking findings of autosomal-dominant cardiomyopathy induced by R120G mutation in the cognate chaperone protein, CRYAB. In both syndromes, we demonstrate increased partitioning of CRYAB phosphorylated on serine-59 to NP40-insoluble aggregate-rich biochemical fraction. While CRYAB undergoes phase separation to form condensates, the phospho-mimetic mutation of serine-59 to aspartate (S59D) in CRYAB mimics R120G-CRYAB mutants with reduced condensate fluidity, formation of protein aggregates and increased cell death. Conversely, changing serine to alanine (phosphorylation-deficient mutation) at position 59 (S59A) restored condensate fluidity, and reduced both R120G-CRYAB aggregates and cell death. In mice, S59D CRYAB knock-in was sufficient to induce desmin mis-localization and myocardial protein aggregates, while S59A CRYAB knock-in rescued left ventricular systolic dysfunction post-myocardial infarction and preserved desmin localization with reduced myocardial protein aggregates. 25-Hydroxycholesterol attenuated CRYAB serine-59 phosphorylation and rescued post-myocardial infarction adverse remodeling. Thus, targeting CRYAB phosphorylation-induced condensatopathy is an attractive strategy to counter ischemic cardiomyopathy.

## Introduction

Cardiomyopathy is characterized by altered structure and function of the cardiac contractile apparatus, whereby targeted approaches to restore sarcomere structure and function hold therapeutic promise. While genetic mutations in contractile proteins reveal mechanisms that have spurred development of myosin modulators to treat hypertrophic cardiomyopathy (1), mechanistic studies targeting the cardiac contractile apparatus in more common causes of cardiomyopathy are lacking. As a case in point, myocardial infarction and ischemia are implicated in the pathogenesis of nearly two-thirds of all cardiomyopathy and heart failure; and abnormal sarcomere function (2) with transcriptional (3) and post-translational modifications of sarcomeric proteins (4, 5) have been described in ischemic cardiomyopathy. Similarly, studies have demonstrated loss of sarcomeres with decompensation of pressure overload hypertrophy to heart failure (6), a common pathophysiologic mechanism implicated in causing heart failure. Given that state-of-the art strategies to treat cardiomyopathy, including myocardial reperfusion, neurohormonal antagonism and resynchronization are only partially effective in improving outcomes (7), there is an urgent unmet need to treat the pathology affecting the cardiac myocyte contractile apparatus in these highly prevalent causes of cardiomyopathy.

Cardiac myocytes rely on intricately coordinated protein quality control mechanisms to maintain structure and function, and drive uninterrupted sarcomere shortening and relaxation that underlies cardiac function (8). These homeostatic mechanisms respond rapidly to stress, regulating various steps over the life cycle of individual proteins. Chaperone proteins play a central role in maintaining cardiac myocyte protein quality, and mutations in CRYAB (HSPB5), HSP27 (HSPB7), HSPB6, HSPB8, and co-chaperones such as BAG3 are implicated in pathogenesis of human cardiomyopathies by affecting stability, localization or turnover of sarcomeric proteins (9).

Mechanisms that drive cardiomyopathy in these rare genetic disorders may offer clues to understanding the molecular basis for sarcomeric abnormalities in more common causes of heart failure, and inform novel approaches for prevention and treatment of associated morbidity and mortality.

Intriguingly, many disease-associated mutations in cardiac chaperone proteins induce intra-cellular protein-aggregate pathology in cardiac myocytes (10–12). Cardiac myocyte protein-aggregate pathology is also observed with human idiopathic dilated cardiomyopathy, ischemic cardiomyopathy and hypertrophic cardiomyopathy (13–16). Desmin, a sarcomere-associated protein, plays a critical role in maintenance of sarcomere structure and subcellular registration; and mutations in desmin or its chaperone CRYAB, result in functional deficiency of desmin or its mis-localization to aggregates, resulting in a group of disorders termed ‘desminopathies’ (17). In some studies, reduced phosphorylation of desmin and its cleavage have also been associated with development of pressure-overload induced cardiomyopathy (16). We and others have demonstrated that the mutant R120G protein binds with increased affinity with desmin to sequester it in aggregates (18, 19); and stimulation of the autophagy-lysosome pathway removes CRYAB-R120G mutant proteins and its aggregates and restores normal desmin localization (18, 20). Notably, this phenomenon of aggregate-prone proteins hijacking normal cardiac myocyte proteins is also observed with expression of P209L mutation of human BAG3, which induces restrictive cardiomyopathy in humans (21). Transgenic expression of P209L BAG3 in the mouse heart induced protein-aggregates, with sequestration of endogenous wild-type BAG3 as well as sarcomere-associated proteins such as desmin, α-actinin and myopodin (SYNPO-2) in aggregates, provoking sarcomere disruption and cardiomyopathy (22). Therefore, examining the mechanisms by which proteins become aggregate-prone holds therapeutic promise to prevent proteotoxic cardiac dysfunction.

To understand the mechanisms by which chaperone-protein interactions are pathologically altered from dynamic protein assemblies in physiology to persistent protein-aggregates in pathology (8), we focused on the phenomenon of phase separation of biomolecules, to form membrane-less compartments termed ‘condensates’ (23). The biophysical principles of phase separation rely on multivalent interactions, realized by oligomerization of ‘intrinsically disordered regions’ (IDRs) within proteins that do not display a fixed tertiary structure, and combine with similar domains in proteins, or interact with other biomolecules (24). Evidence is accumulating that mutations or protein modifications cause biomolecular condensates to transition from a ‘liquid state’ to a ‘gel-like state’ to nucleate conversion to protein-aggregates that are observed in a variety of disease states (25, 26). We discovered that CRYAB undergoes phase separation and stress-induced phosphorylation at serine-59, which is also increased in the aggregate-prone human-disease causing R120G mutant. This alters ‘liquid-like’ properties of CRYAB condensates towards ‘gel-like’ behavior with reduced fluidity, and aggregate formation conferring cellular toxicity. Treatment with 25-hydroxycholesterol, a compound that ameliorates CRYAB-R120G aggregates in the lens (27), limited stress-induced phosphorylation of CRYAB and the resultant improvement in condensate properties was associated with attenuation of post-myocardial infarction (post-MI) ventricular remodeling.

Taken together, our data demonstrate a mechanistic role of altered condensate behavior of CRYAB, i.e. a ‘condensatopathy (28)’ in the pathogenesis of ischemic cardiomyopathy and support the development of pharmacologic approaches to reduce the abundance of this aggregate-prone toxic protein to treat this common cause of heart failure.

## Results

### Desmin mis-localizes to protein-aggregates in human ischemic cardiomyopathy

Mis-localization of desmin to protein-aggregates is observed with heritable mutations in *DES*, the gene coding for desmin (17, 29) and in *CRYAB* (10), the gene coding for its chaperone protein, αB-crystallin. In these disorders, loss of physiologic desmin localization and consequently its function induces cardiomyopathy that mimics the pathology observed with experimental mouse ablation of the *DES* gene in mice (30), whereby these disorders are termed ‘desminopathies’ (17). To examine desmin localization in ischemic cardiomyopathy (ICM), we performed immunohistochemistry and biochemical subcellular fractionation on human heart tissue obtained from humans with ICM and from donor hearts without known cardiac pathology, that were not used for transplantation (Table S1). As shown in Fig. 1A, desmin was observed along the Z-discs and intercalated discs in donor hearts consistent with its physiologic localization. In contrast, desmin was mis-localized (with reduced localization in striations assessed as worse striation score, see legend) to protein-aggregates in ICM hearts (Fig. 1A, B) along with polyubiquitinated (polyUb) proteins (Fig. S1A). This was accompanied by immunodetectable pre-amyloid oligomers in ICM hearts as detected by A11 antibody (15) staining (Fig. S1B). Importantly, we also detected mis-localization of actin, another CRYAB client protein (31) as well as of α-actinin (a Z-disc protein that is observed to interact with CRYAB (32)); with increased localization of these proteins to aggregates in ICM hearts (Fig. 1A, B). This was accompanied by increased abundance of p62 (an adaptor protein that binds to and sequesters polyUb proteins in aggregates (33, 34)) (Fig. 1C, D), and increased polyUb proteins (Fig. 1C, E) in the aggregate-rich NP40 detergent-insoluble fraction suggesting sequestration of cardiac myocyte proteins in aggregates in ICM hearts.

**Figure 1:**
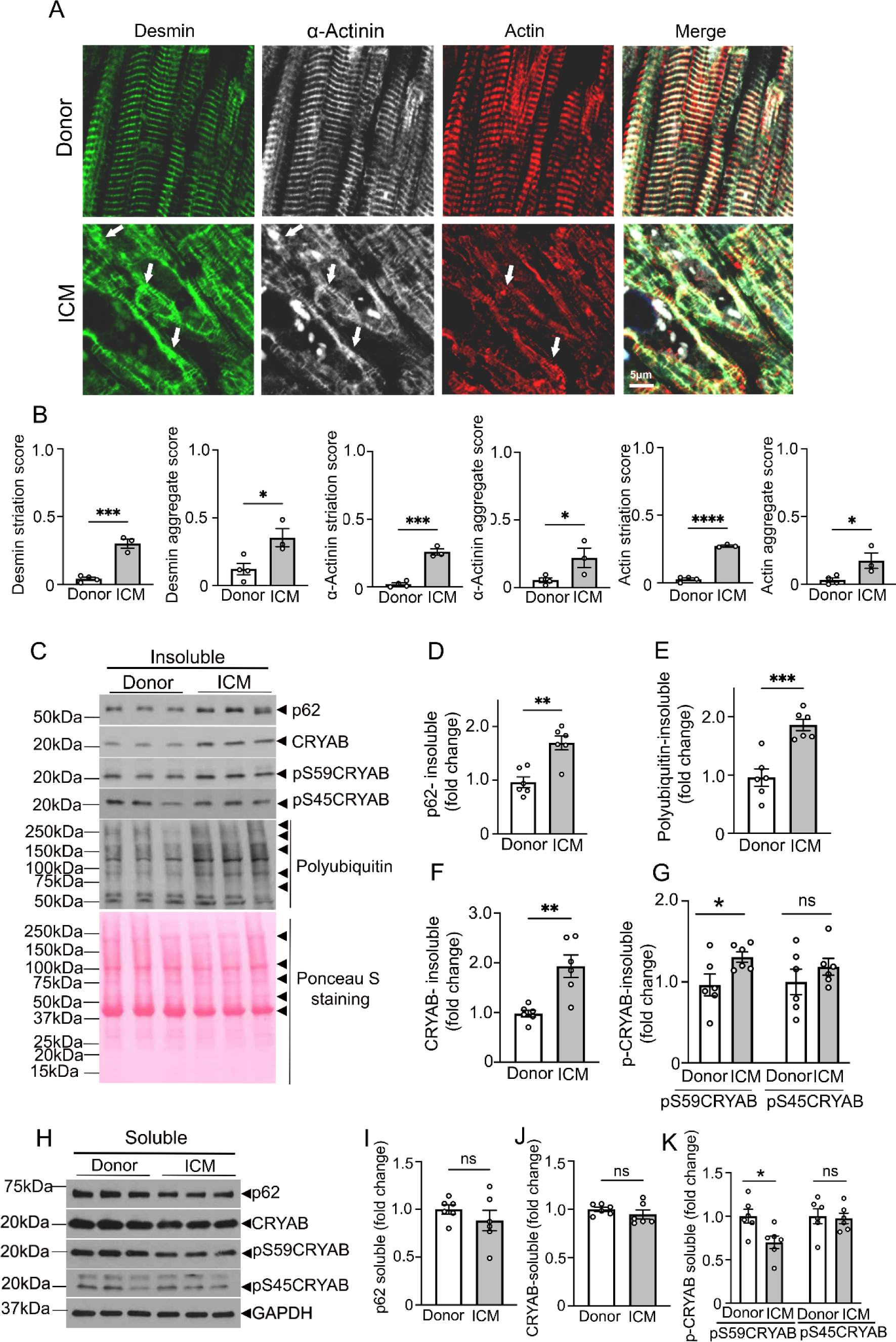
Desmin, α-Actinin and Actin, and the serine-59 phosphorylated form of their chaperone protein, CRYAB, localize to protein-aggregates in human ischemic cardiomyopathy. A. Representative immunohistochemical images from left ventricular myocardium of individuals evaluated as controls (donor) or patients with end-stage ischemic cardiomyopathy (ICM) stained for desmin, α-actinin and actin. Arrows point to mis-localization of these proteins from their physiologic location on Z-discs and intercalated discs (desmin), Z-disc (α-actinin) and sarcomere (actin) in donor myocardium to protein-aggregates in ICM myocardium. B. Quantitation of striation score and aggregate score for desmin, α-actinin and actin in ICM and donor hearts. N=3-4 hearts/group. For striation scoring, normal localization of proteins got scored as 0; and abnormal striation or mis-localization of proteins was scored as of 1. For scoring aggregates, absence of aggregates was scored as 0 and presence of aggregates was scored as 2. C-G. Immunoblot (C) and quantitation (fold change as compared to donor mean) depicting total p62 (D), polyubiquitinated proteins (E), CRYAB (F) and pS59-CRYAB and pS45CRYAB (G) in NP40-detergent-insoluble fractions from human hearts from patients with ischemic cardiomyopathy (ICM) and donors. Ponceau S staining is shown as loading control. H-K. Immunoblot (H) and quantitation for p62 (I), CRYAB (J), and pS59-CRYAB and pS45CRYAB (K) abundance in NP-40 detergent soluble biochemical fractions from human hearts as in C-G. GAPDH was used as loading control. N=6 samples/group for C-K. * indicates P <0.05, ** indicates P<0.01 and *** indicates P<0.001 vs. donor as control by t-test. ‘ns’ indicates not significant.

To examine potential mechanisms for mis-localization of sarcomeric (actin, α-actinin) and sarcomere-associated proteins (desmin), we evaluated the localization of CRYAB, which plays an important role in chaperoning these proteins (19). We observed increased partitioning of CRYAB to the detergent-insoluble aggregate-rich fraction (Fig. 1C, F)). Interestingly, prior studies indicate that CRYAB undergoes post-translational modifications, specifically phosphorylation at serine resides (at position 19, 45 and 59), under stress (35). Of these, serine 45 and 59 are phosphorylated by stress-induced p38 mitogen-activated protein kinase (MAPK) (35), as well as by protein kinase N, which are activated in cardiac ischemia-reperfusion injury (36, 37). Also, studies in cell-culture models show that aggregate-prone R120G mutant of CRYAB is hyperphosphorylated at these three serine residues, which regulates its tendency to aggregate (38). Our findings demonstrate increased abundance of phosphorylated CRYAB at residue serine-59 (pS59-CRYAB), but not at serine 45 in the detergent-insoluble fraction (Fig. 1C, G) with a concomitant decline in its abundance in the soluble fraction (Fig. 1H, K), which parallels the partitioning of p62 and polyUb proteins to the aggregate-rich detergent-insoluble fraction.

To examine if partitioning of pS59-CRYAB is also observed in mouse myocardium under stress, we performed closed-chest cardiac ischemia-reperfusion (IR) injury in young adult wild-type C57BL/6J mice and evaluated them 4 weeks later. As shown in Supplemental Figure S2A, B; IR injury induced a marked increase in left ventricular end-diastolic volume (EDV) with decline in left ventricular ejection fraction (EF) at 4 weeks post-MI, as compared with sham operated-mice, consistent with development of ischemic cardiomyopathy. Similar to our observations in human ICM (Fig. 1), this was accompanied by a significant increase in abundance of pS59-CRYAB with increased p62 in the aggregate-rich NP40-insoluble fraction (Figure S2C-I) and mis-localization of desmin, α-actinin and actin in the post-IR myocardium (Fig. S2J, K). Taken together, these findings demonstrate that CRYAB is increasingly phosphorylated at serine-59 in the setting of ischemic cardiomyopathy in mice and in humans, and partitions into aggregate-rich biochemical protein fraction. This associated mis-localization of desmin (and other CRYAB clients) to protein-aggregates suggests the hypothesis that phosphorylation at serine-59 may render CRYAB aggregate-prone and sequester its client proteins within protein-aggregates in ischemic cardiomyopathy. Indeed, the aggregate-prone CRYAB-R120G mutant protein is phosphorylated at serine-59 in the mouse myocardium in transgenic R120G mice (Fig. S3), confirming prior observations in cell culture (38), in a mouse model for proteotoxic cardiomyopathy that recapitulates human pathology with mis-localization of desmin to protein aggregates (18) to induce a ‘desminopathy’. Taken together with our previous observation of increased affinity for R120G mutant protein to bind to desmin as compared with native CRYAB (18), these findings suggest the hypothesis that increased pS59-CRYAB binds to its client proteins with increased affinity to sequester them in aggregates in ischemic cardiomyopathy.

### Phosphorylation at serine-59 is necessary and sufficient to make CRYAB aggregate-prone

To examine the functional relevance of serine-59 phosphorylation, we generated CRYAB mutants that mimic a phosphorylation-deficient state at that residue by replacing serine with alanine (S59A) or a phospho-mimetic state with a change to aspartic acid (S59D). We also generated the CRYAB- R120G mutant with the S59A change and expressed these mutants with a N-terminal GFP tag in HEK293 cells to examine the relevance of serine-59 phosphorylation in regulating its aggregation potential. Despite being expressed at equivalent levels (Fig. 2A), the S59D change resulted in formation of GFP-positive CRYAB aggregates mimicking the observations with the R120G mutant (Fig. 2B, C). By contrast, the S59A change markedly reduced the aggregation of the CRYAB-R120G mutant protein indicating that serine-59 phosphorylation is necessary for its aggregate-prone behavior (Fig. 2B, C). As we have previously demonstrated (18), aggregate-prone CRYAB-R120G was toxic and induced cell death (Fig. 2D). The S59D mutant was indeed sufficient to induce increased cytotoxicity as compared with wild-type CRYAB, whereas the S59A mutant attenuated the toxicity of the CRYAB-R120G mutant (to a similar extent as observed with CRYAB triple mutant with serine to alanine change at 19, 45 and 59; Fig. S4A-C), paralleling the observations with their aggregate-prone behavior (Fig. 2D).

**Figure 2:**
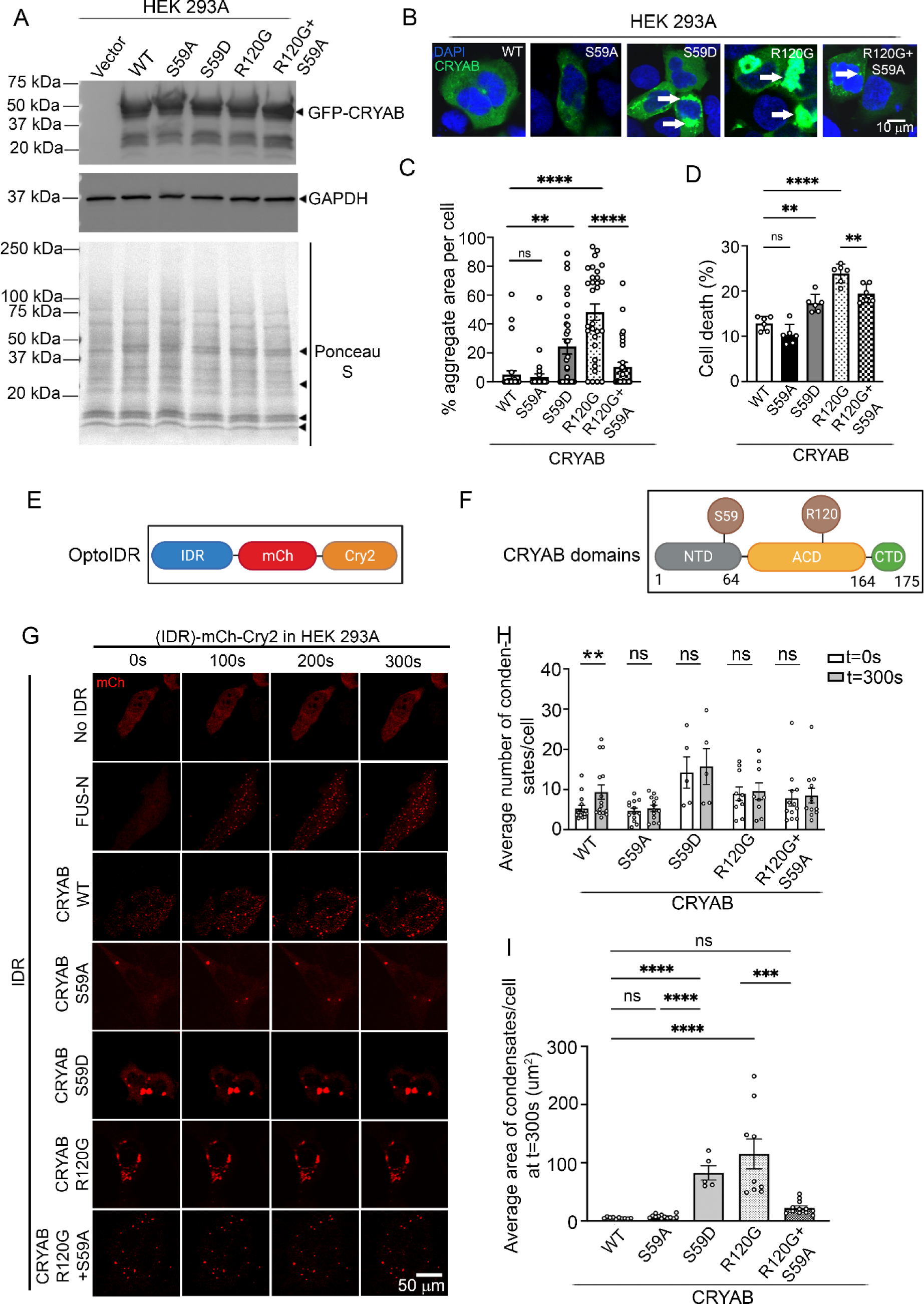
Phosphorylation of CRYAB at S59 makes it aggregate-prone and toxic. **A**) Immunoblot (A) demonstrating expression of GFP-fusion proteins in HEK293A cells transfected with GFP-tagged wild-type CRYAB, its phospho-mimetic mutant (S59D), phosphorylation-deficient mutant (S59A), R120G mutant or the R120G and S59A double mutant proteins. **B, C)** Representative immunofluorescence images (B) for detection of protein-aggregates with quantitation (C) of aggregate area per cell. ** indicates P<0.01, *** indicates P<0.001 and **** indicates P<0.0001 by Tukey’s post-hoc test after one-way ANOVA. Nuclei are blue (DAPI). **D)** Cell death in cells treated in A. *** indicates P<0.001 and **** indicates P<0.0001 by Tukey’s post-hoc test after one-way ANOVA. **E**) Schematic depicting generation of optoIDR constructs. ‘IDR’ indicates intrinsically disordered region, mCh indicates mCherry fluorophone and Cry2 encodes for *Arabidopsis thaliana* protein with light-activated phase separation characteristics. **F)** Various domains of CRYAB with localization of serine-59 and arginine 120 residues depicted. **G)** Representative time-lapse images at t=0s, 100s, 200s, and 300s after light activation in HEK293A cells transfected with constructs generated with CRYAB WT, its phosphorylation-deficient mutant (S59A), phospho-mimetic mutant (S59D), R120G mutant or the R120G and S59A double mutant proteins as the ‘IDR’ in the optoIDR constructs. Cry2 fused with mCherry without an IDR was used as the negative control and FUS-N fused with mCherry-Cry2 was studied as positive control. **H)** Average number of condensates/cell at t=0 vs. t=300s in cells treated as in E. ** indicates P<0.01 by Mann-Whitney test. **I)** Average area of condensates/cell at t=300s in cells treated in E. *** indicates P<0.001 and **** indicates P<0.0001 by Tukey’s post-hoc test after one-way ANOVA. ‘ns’ indicates not significant for all panels.

### Phosphorylation at serine-59 alters the phase separation behavior of CRYAB

An emerging body of evidence indicates that phase separation of proteins regulates their ability to form biomolecular assemblies termed as ‘condensates’, and the biophysical properties that determine the dynamicity and fluidity of condensates regulate their aggregation potential (24, 26, 28). Accordingly, to examine whether CRYAB can phase separate in living cells, we adapted the OptoDroplet assay system (39) and expressed full length CRYAB, as well as its N-terminus, C-terminus and α-crystallin domain (ACD) separately (Fig. 2E, F). We employed the N-terminal domain of FUS (a condensate forming protein implicated in neurodegeneration as a positive control) and a Cry2 construct lacking an IDR as a negative control (39) and examined their propensity to phase separate after light activation. As shown, light activation resulted in dynamic phase separation of CRYAB into condensates mimicking the observations with N-terminal fragment of FUS protein, while Cry2 protein by itself did not phase separate, as previously described (39) (Fig. 2G, video S1, S2). Remarkably, CRYAB full-length protein and each of CRYAB’s N-terminus, C-terminus and α-crystallin domain (ACD) domains demonstrate the propensity to phase separate (Fig 2G, H; Fig. S5, Video S3, S4, S5, S6). While some spherical CRYAB condensates were observed at baseline, prior to light activation, the average number of condensates doubled after light activation (Fig. 2G, H, video S3). Interestingly, a S59A or S59D change in the full-length protein completely abrogated light-induced condensate formation (Fig. 2G, H; video S7, S8). However, like the observations with GFP-tagged proteins (Fig. 2A-C), the mCherry-tagged optoIDR constructs induced protein-aggregates in cells expressing S59D and CRYAB-R120G mutant proteins (Fig. 2G-I; Video S8, S9) even prior to light activation, with markedly larger and irregularly shaped protein-aggregates in S59D transfected cells mimicking the R120G mutant (Fig. 2G, I). In agreement with the observations with the GFP-tagged CRYAB- R120G-S59A vs. the CRYAB-R120G constructs (Fig. 2A-C), the S59A change reduced the size of R120G optoDroplet construct aggregates (Fig. 2G, I; Video S10); but did not result in a noticeable light-induced increase in condensate formation (Fig. 2G, H).

These findings suggest that similar to the R120G mutation, the S59D change reduces the fluidity of condensates, making CRYAB aggregate prone. To examine this premise, we performed fluorescence recovery after photobleaching (FRAP). As shown in Fig. 3A-C, WT CRYAB shows rapid recovery following photobleaching, indicating that these phase-separated condensates are dynamic and liquid-like. In contrast, the recovery of fluorescence was markedly reduced in S59D and R120G mutant proteins as compared with wild-type CRYAB (Fig. 3A-C). Remarkably, the S59A change in R120G restored its fluorescent recovery to wild-type levels (Fig. 3A-C). The fluorescence recovery was comparable in the S59A mutant to wild-type CRYAB (Fig. 3A-C). Taken together, these data indicate that serine-59 phosphorylation is both necessary and sufficient to alter the phase separation behavior of CRYAB, reducing fluidity of CRYAB condensates to convert them to a ‘gel-like’ state (25, 26), as a potential explanation for formation of aggregates.

**Figure 3:**
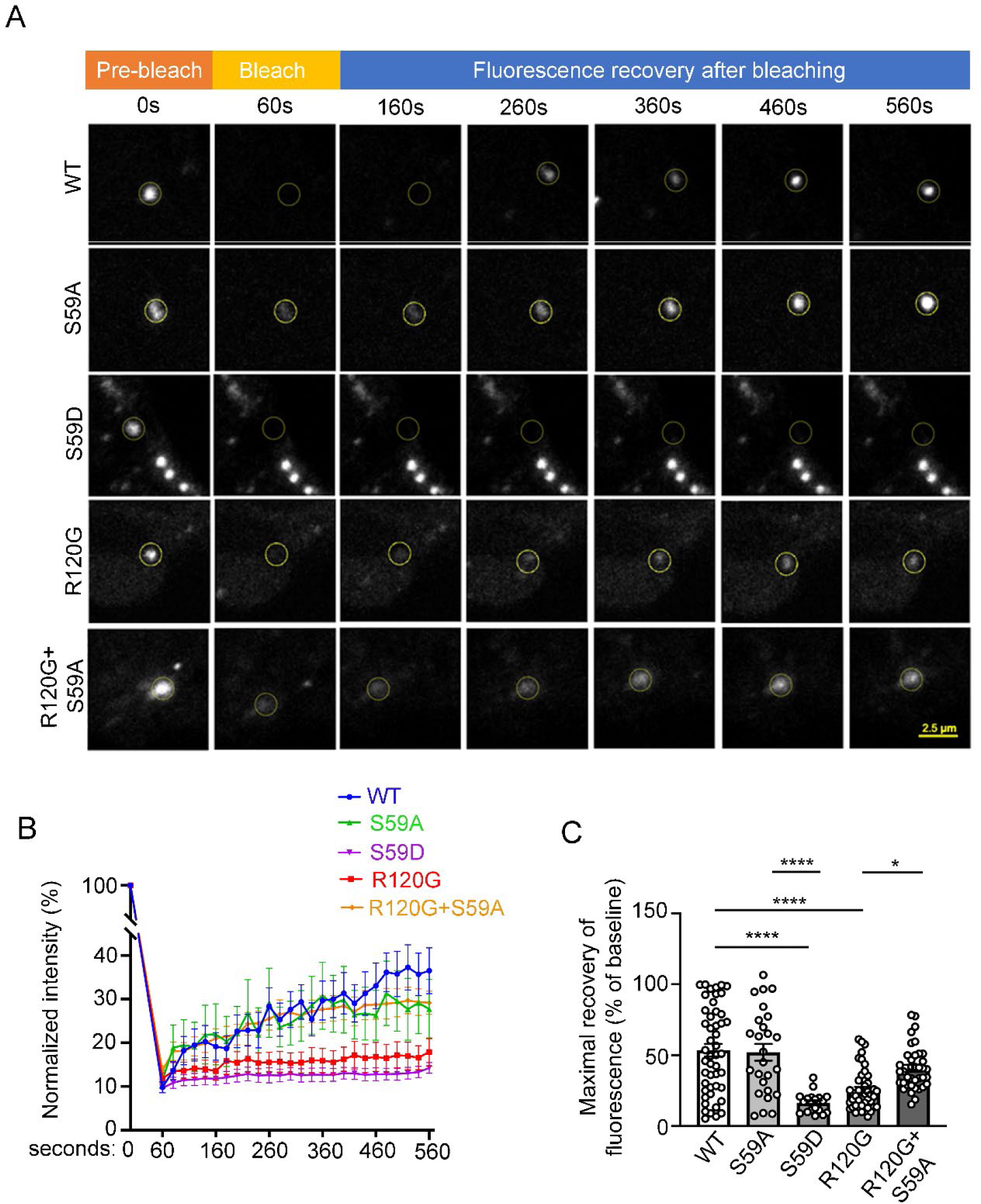
Phosphorylation of CRYAB at serine-59 reduces dynamicity of condensates. **A, B)** Representative images demonstrating recovery of fluorescence after photobleaching in HEK 293A cells transfected with mCherry-Cry2 fused optoIDR constructs generated with CRYAB WT, its phosphorylation-deficient mutant (S59A), phospho-mimetic mutant (S59D), R120G mutant, or the R120G and S59A double mutant proteins. Representative images demonstrate area of photobleaching (marked with a dotted circle) prior to (pre-bleach), immediately after, and at 100, 200, 300, 400 and 500 seconds (s) after photobleaching was terminated. Intensity at various time points is depicted as a fraction of intensity prior to bleaching (set at 100%). **B, C.** Fluorescence intensity normalized to baseline (B) and quantitation of fluorescence recovery (C, maximum minus immediately post-bleach) in condensates of various CRYAB variants indicated in A. * indicates P<0.05, and **** indicates P<0.0001 by Tukey’s post-hoc test after one-way ANOVA.

### Mice bearing phospho-mimetic aspartic acid residue in CRYAB at position 59 demonstrate protein aggregates in the myocardium

To rigorously determine the role of phosphorylation of the serine-59 residue of CRYAB in myocardial homeostasis and response to stress, we used CRISPR-Cas9 editing to generate mice homozygous for phosphorylation deficient serine to alanine change (S59A) and phospho-mimetic serine to aspartic acid change (S59D). We confirmed the loss of immuno-detectable pS59 CRYAB in the S59A myocardium (Fig. S6). Either amino acid substitution did not alter total CRYB levels (Fig. S7), left ventricular structure and function or body weight (Table S2) or result in myocardial histologic abnormalities in young adult mice of both genotypes (Fig. 4A); which were also grossly indistinguishable from wild type mice. However, transmission electron microscopy revealed abnormal mitochondria with cristal rarefaction and presence of protein aggregates in cardiac myocytes from S59D mice (Fig. 4A). The myocardium in S59A mice was ultra-structurally indistinguishable from those of wild type mice (Fig. 4A). This was correlated with biochemical evidence of increased polyubiquitinated proteins in the aggregate-rich insoluble fraction (Fig. 4C, D) and increased p62 in both soluble and insoluble fractions (Fig. 4C, E) in myocardium from S59D mice as compared with wild type and S59A mice. Notably, the there was a reduction in CRYAB protein abundance in the soluble fraction from S59D mice and a concomitant increase in the aggregate-rich insoluble fraction as compared with the other two genotypes (Fig. 4B, C, F), pointing to increased propensity of S59D CRYAB to form protein aggregates as we have observed previously (Fig. 2A-C). Mice bearing phosphorylation-deficient serine to alanine change in CRYAB were indistinguishable from wild type on the above parameters.

**Figure 4:**
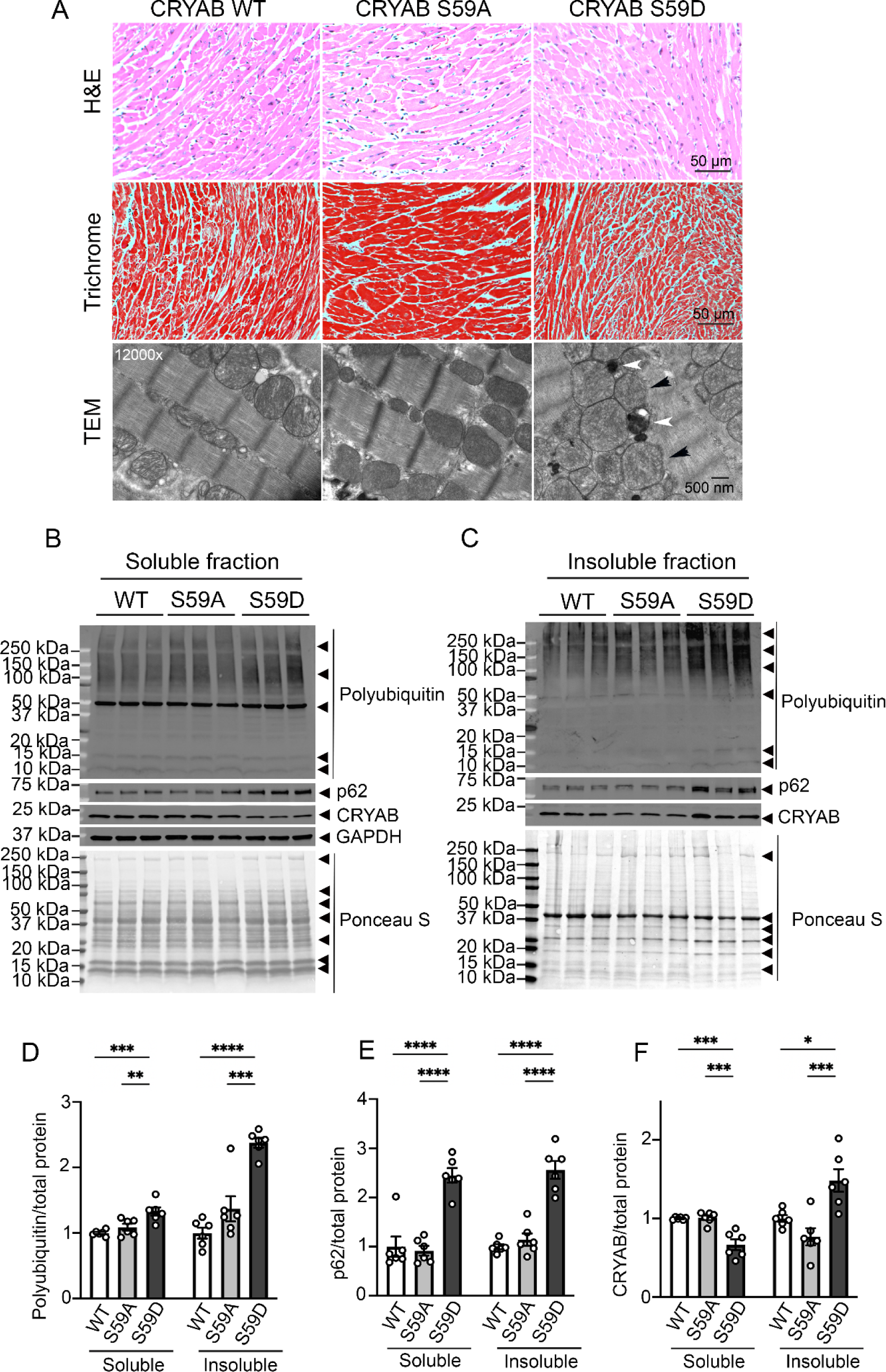
Mice with phospho-mimetic aspartic acid residue instead of serine at position 59 in CRYAB demonstrate myocardial protein aggregates. **A)** Representative hematoxylin-eosin (*top row*) and Masson’s trichrome-stained (*middle row*) myocardial sections from young adult mice homozygous for alleles bearing serine to aspartic acid (phospho-mimetic residue; S59D) or serine to alanine (phosphorylation-deficient; S59A) mutation at position 59 in CRYAB; and mice bearing wild-type (WT) CRYAB alleles as controls. Transmission electron micrographs (*bottom row*) from wild type, S59A and S59D mice. Representative of n=2 mice per group. Black arrows point to abnormal appearing mitochondria and white arrows point to protein aggregates. **B, C)** Representative immunoblots depicting expression of poly-ubiquitinated proteins and p62, CRYAB and GAPDH proteins in NP40-soluble (B) and NP40-insoluble (C) myocardial extracts from wild type, S59A and S59D mice. Ponceau S staining is shown as loading control. **D-F)** Quantitation of poly-ubiquitinated proteins (D), p62 (E) and CRYAB (F) in soluble and insoluble myocardial extracts from wild type, S59A and S59D mice. * indicates P<0.05, ** indicates P<0.01, *** indicates P<0.001 and **** indicates P<0.0001 by Tukey’s post-hoc test after one-way ANOVA.

### Phosphorylation-deficient serine to alanine change in CRYAB at position 59 attenuates post-myocardial infarction left ventricular systolic dysfunction

To test the hypothesis that stress-induced phosphorylation of serine 59 in CRYAB, in the post-myocardial infarction myocardium is a mechanism for adverse left ventricular remodeling (as observed in Fig. S2), we subjected young adult S59A and S59D mice and wild type controls to closed chest ischemia reperfusion (IR) modeling and examined left ventricular structure, function and scar size at 4 weeks post-MI (Fig. 5A). Mice bearing S59A CRYAB alleles demonstrated improved left ventricular ejection fraction at 4 weeks post-MI as compared with wild type mice and with mice bearing S59D alleles (Fig. 5D) despite comparable extent of injury (as determined by area-at-risk, Fig. 5B). Also, left ventricular dilation post-MI was significantly reduced in S59A versus S59D mice (Fig. 5C) indicating that preventing phosphorylation at serine 59 abrogated adverse left ventricular remodeling post-MI. This was also accompanied by reduced scar size in the S59A mice as compared with the other two genotypes (Fig. 5E, F). Taken together, these data indicate that phosphorylation as serine 59 contributes to adverse left ventricular remodeling post-myocardial infarction.

**Figure 5:**
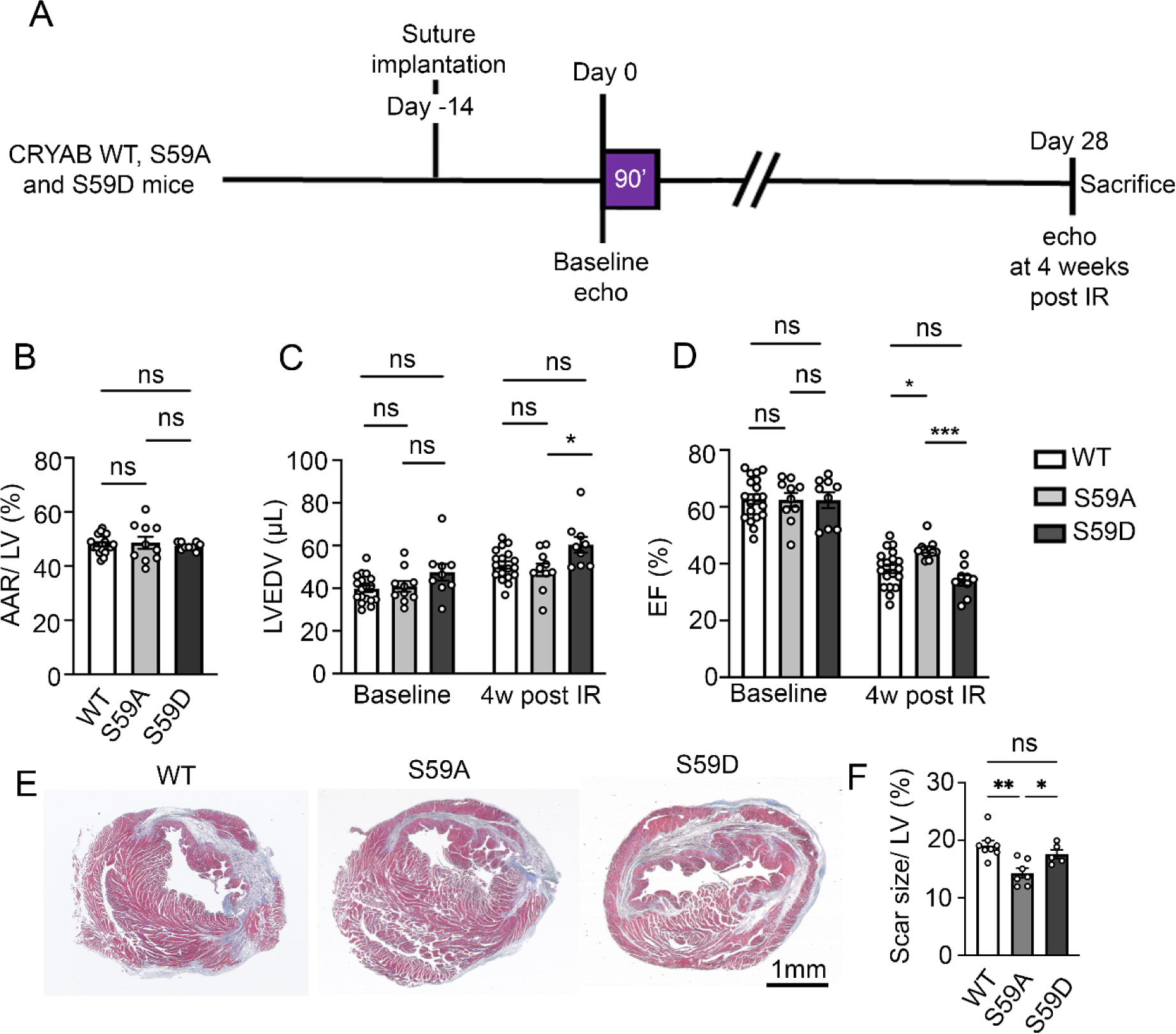
Knock-in of phosphorylation-deficient alanine instead of serine at position 59 in CRYAB rescues post-IR LV remodeling in mice. **A)** Schematic depicting experimental strategy for closed-chest IR modeling (90 minutes of ischemia followed by reperfusion) in S59A, S59D and wild type (WT mice). **B)** Quantitative assessment of area-at-risk (AAR) during LAD occlusion in mice treated as in A. **C, D)** Quantitative analyses of left ventricular end-diastolic volume (LVEDV, C) and LV ejection fraction (EF (%), D) at baseline (i.e. prior to) and at 4 weeks post-IR injury. * denotes P < 0.05, ** denotes P <0.01 and *** denotes P <0.001 by Tukey’s post-hoc testing after one-way ANOVA. Echocardiographic parameters at baseline and 4 weeks post-IR injury are analyzed separately as they were performed under different anesthetic regimens. **E, F)** Masson’s trichrome stained left ventricular sections demonstrating presence of scar at 4 weeks post-IR injury (E) with quantitation of scar size (F). * denotes P <0.05 and ** denotes P <0.01 by Tukey’s post-hoc testing after one-way ANOVA.

### Phosphorylation of CRYAB at serine 59 increases its interaction with desmin resulting in its mis-localization to protein aggregates

We next examined the effect of CRYAB phosphorylation on its interaction with its client protein desmin in the myocardium using mice modeled to generate phosphorylation deficient (S59A) and phospho-mimetic (S59D) change in CRYAB. Desmin was mis-localized from striations with increased aggregates at 4 weeks post ischemia-reperfusion injury (IR) with increase in poly- ubiquitinated proteins in the myocardium as compared with sham controls (Fig. 6A-C).

**Figure 6:**
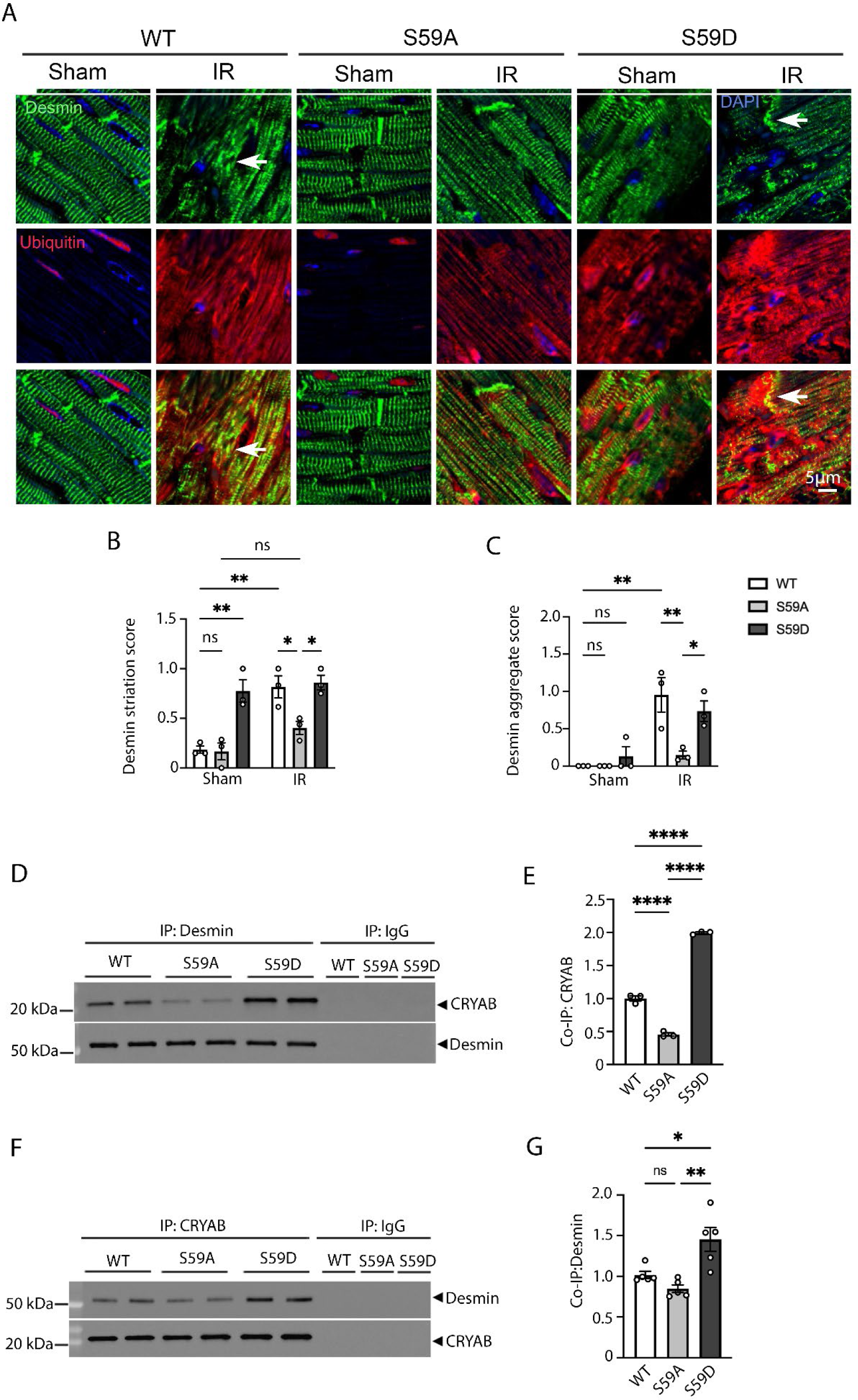
Knock-in of phosphorylation-deficient alanine or phospho-mimetic aspartic acid instead of serine at position 59 in CRYAB have opposing effects on desmin localization. **A)** Representative immuno-histochemical images demonstrating localization of desmin (*top row*) and polyubiquitinated proteins (*middle row*) and merged images (*bottom row*) from S59A, S59D and wild-type (WT) mice 4 weeks after being subjected to closed chest IR injury or sham procedure. Arrows point to desmin aggregates. **B, C)** Quantitative evaluation of desmin localization with striation score (B) and aggregated desmin (C) in S59A, S59D and wild-type (WT) mice 4 weeks after being subjected to closed chest IR injury or sham procedure. For striation scoring, normal localization of proteins got scored as 0; and abnormal striation or mis-localization of proteins was scored as of 1. For scoring aggregates, absence of aggregates was scored as 0 and presence of aggregates was scored as 2. * denotes P < 0.05 and ** denotes P <0.01 by Tukey’s post-hoc testing after one-way ANOVA. **D, E)** Representative immunoblot (D) and quantitative assessment (E) of CRYAB and desmin expression after immuno-precipitation (IP) with desmin from myocardial extracts from young adult S59A, S59D and wild-type (WT) mice. Expression of CRYAB is assessed as fold of WT control. **F, G)** Representative immunoblot (F) and quantitative assessment (G) of CRYAB and desmin expression after immuno-precipitation (IP) with CRYAB from myocardial extracts from young adult S59A, S59D and wild-type (WT) mice. Expression of desmin is assessed as fold of WT control. * denotes P < 0.05, ** denotes P <0.01, *** denotes P <0.001 and **** denotes P <0.0001 by Tukey’s post-hoc testing after one-way ANOVA.

Interestingly, as compared with wild type in the post-MI myocardium, normal desmin localization to striations was better preserved in S59A with reduced desmin aggregates (Fig. 6A-C). By contrast, desmin was mis-localized with increased aggregates even in sham treated S59D myocardium and this persisted at 4 weeks post-MI (Fig. 6A-C). A potential explanation for these findings is the increased interaction between CRYAB phosphorylated at serine 59 with desmin. Indeed, we have previously observed that R120G mutant CRYAB (which is predominantly phosphorylated at the serine 59 residue, Fig. S3) demonstrates increased interaction with desmin as compared with wild-type CRYAB (18). Accordingly, we performed immuno-precipitation (IP) studies to examine the interaction between CRYAB and desmin in the myocardium from unstressed and young adult S59A, S59D and wild type mice, and observed increased interaction between desmin and S59D as compared with wild type CRYAB and reduced interaction between desmin and S59A CRYAB as compared with wild-type CRYAB (Fig. 6-D-G); a finding that was confirmed by reverse co-IP studies as shown. Taken together, these findings suggest that phosphorylation of CRYAB at serine 59 alters its condensate properties to make it aggregate prone and increases its binding with client proteins as the mechanism for mis-localization of these proteins (such desmin, α-actinin and actin as in Fig. 1A) to aggregates.

### Attenuating serine-59 phosphorylation with 25-hydroxycholesterol rescues post-IR cardiomyopathy

Our data indicate that pS59-CRYAB is a toxic aggregate-prone protein, and a phosphorylation deficient mutant of CRYAB attenuates post-MI adverse left ventricular remodeling. To examine the paradigm that a pharmacologically induced reduction in pS59-CRYAB may be beneficial in post-MI setting, we turned to studies with 25-hydroxycholesterol (25-HC). This compound was uncovered in a screen focused on preventing aggregation of CRYAB-R120G in the lens of the eye to prevent cataract formation (27). It has been predicted to bind in a groove in CRYAB (40), whereby we hypothesized that it will prevent phosphorylation of CRYAB, perhaps by restricting access to serine 59. Accordingly, we tested the efficacy of 25-HC on serine-59 phosphorylation of CRYAB-R120G protein transduced in neonatal rat cardiac myocytes, followed by subcellular fractionation. As shown, 25-HC induced a dose-dependent reduction in pS59-CRYAB levels in the detergent insoluble fraction (Fig. 7B) with a significant reduction at the highest dose tested (Fig. 7B, 7D) without a change its levels in the soluble fraction (Fig. 7A, D) or in total CRYAB abundance (Fig. 7A, B, E). This was accompanied by a reduction in p62 and polyUb proteins in the insoluble fraction (Fig. 7B, F, G) but not the soluble fraction (Fig. 7A, F, G) with 40µM 25- HC. 25-HC also induced a marked reduction in CRYAB-R120G aggregates detected with immunostaining (Fig. 7C). To determine if 25–HC treatment alters phase separation of the CRYAB-R120G mutant protein, we expressed CRYAB-R120G optoIDR construct (as in Fig. 2E-G) and treated the cells with 40μM 25HC or diluent. 25-HC treatment reduced the number and size of CRYAB-R120G aggregates (Fig. 7H, J), but did not result in a noticeable light-induced increase in condensate formation (Fig. 7H, I, videos S11, S12). To examine the effect of 25-HC treatment on the fluidity of CRYAB-R120G condensates, we performed FRAP analyses. As shown (Fig. 7K, L), the recovery of fluorescence was markedly increased in 25-HC-treated CRYAB- R120G mutant protein post-bleach as compared with diluent-treated CRYAB-R120G. Importantly. 25-HC did not affect condensate number or size, or the fluidity of S59D CRYAB condensates (Fig. S8, videos S13, S14). Taken together, these data indicate that 25-HC treatment reduces serine-59 phosphorylation to maintain fluidity of the resulting condensates and reduces the propensity for aggregate formation.

**Figure 7:**
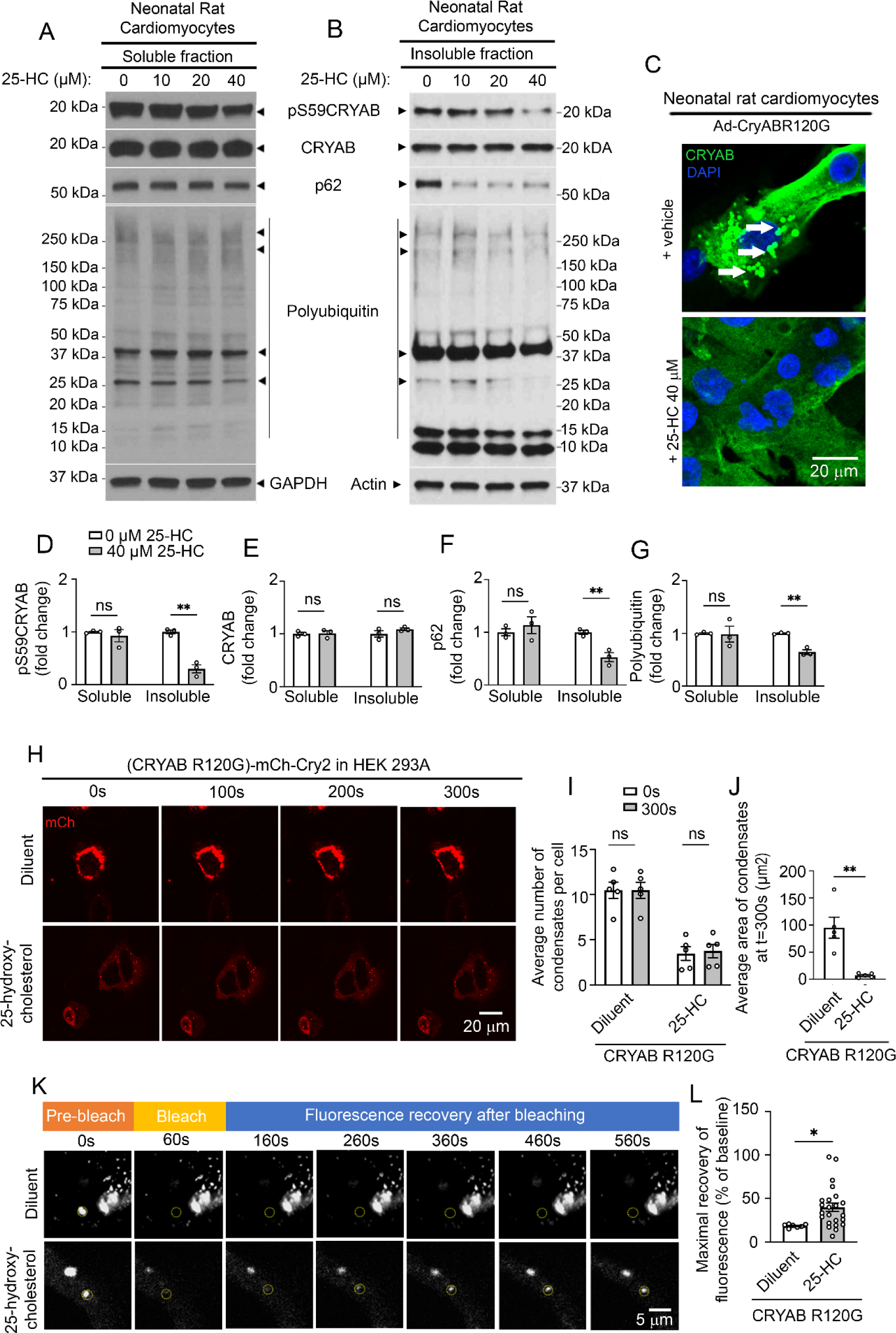
Treatment with 25-hydroxycholesterol reduces phosphorylation of CRYAB at S59 and alters the phase separation characteristics of CRYAB-R120G. **A, B)** Representative immunoblots depicting total pS59-CRYAB, CRYAB, p62 and polyubiquitinated proteins in NP40-detergent (A) soluble and (B) insoluble fractions from neonatal rat cardiomyocytes (NRCMs) transduced with adenoviral CRYAB-R120G (MOI=10) for 90 hours and treated with 0, 10, 20 and 40 mM 25-hydroxycholesterol for the final 72 hours. GAPDH and ACTIN are shown as loading controls. **C.** Representative immunofluorescence images of NRCMs treated in A. Arrows point to GFP-positive aggregates. **D-G)** Quantitative assessment of pS59-CRYAB (D), total CRYAB (E), p62 (F) and polyubiquitinated protein (G) expression in NP-40 soluble and insoluble fractions from NRCMs treated in A. * denotes P <0.05 and ** denotes P <0.01 by t-test. **H)** Representative time-lapse images at t=0s, 100s, 200s, and 300s after light activation in HEK293A cells transfected with OptoIDR constructs generated with CRYAB-R120G as the ‘IDR’ protein and treated with diluent or 25-HC (40mM). **I, J**) Average number (I) and area (J) of condensates/cell at t=0 vs. t=300s in cells treated as in H. *** indicates P<0.001 and **** indicates P<0.0001 by t-test. **K)** Representative images demonstrating recovery of fluorescence after photobleaching in HEK 293A cells treated as in H. The area of photobleaching is marked with a dotted circle prior to (pre-bleach), and at 0, 100, 200, 300, 400 and 500 seconds (s) after photobleaching. **L.** Quantitation of fluorescence recovery in condensates of CRYAB variants (as maximum minus immediately after photobleaching) as shown in K. * indicates P<0.05 by t-test.

These findings predict a beneficial effect of 25-HC on post-IR remodeling by reducing the abundance of pS59-CRYAB. Accordingly, we set up a protocol, where mice were injected with 25-HC at the dose of 10 mg/kg beginning at day 4 after IR injury (Fig. 8A). We chose this time point to avoid its effects on CRYAB phosphorylation during the acute phase of injury, where p38 MAP kinase-mediated serine-59 phosphorylation may play a protective role (41, 42). Our data demonstrate a significant increase in LVEF (Fig. 8D) with reduced LV dilation (Fig. 8C) in 25- HC-treated mice, while the area-at-risk was similar between the two groups (Fig. 8B). We examined the effect of 25-HC on pS59-CRYAB and detected a significant decline in total pS59- CRYAB abundance (Fig. S9). This was accompanied by a decline in pS59-CRYAB levels in the insoluble fraction in mice treated with 25-HC (by 56%, Fig. 8E, G) without a change in the soluble fraction (Fig. 8E, G), consistent with the effects observed in neonatal rat cardiac myocytes (Fig. 7A, B, D). Notably, 25-HC treatment also induced a decline in polyubiquitinated proteins but not p62 in the insoluble fraction (Fig. 8E, I, J). 25-HC-treatment also restored physiologic desmin localization and attenuated aggregates observed in the post-IR myocardium of wild-type mice (Fig. 8K, L, M). There was no significant difference in scar size in 25-HC vs. diluent treated mice post-MI (Fig. 8N, O). These data support the notion that attenuating overall levels of serine-59 phosphorylated CRAYB is an effective strategy to reduce the aggregation potential and toxicity of CRYAB under sustained stress, as observed in ischemic cardiomyopathy.

**Figure 8:**
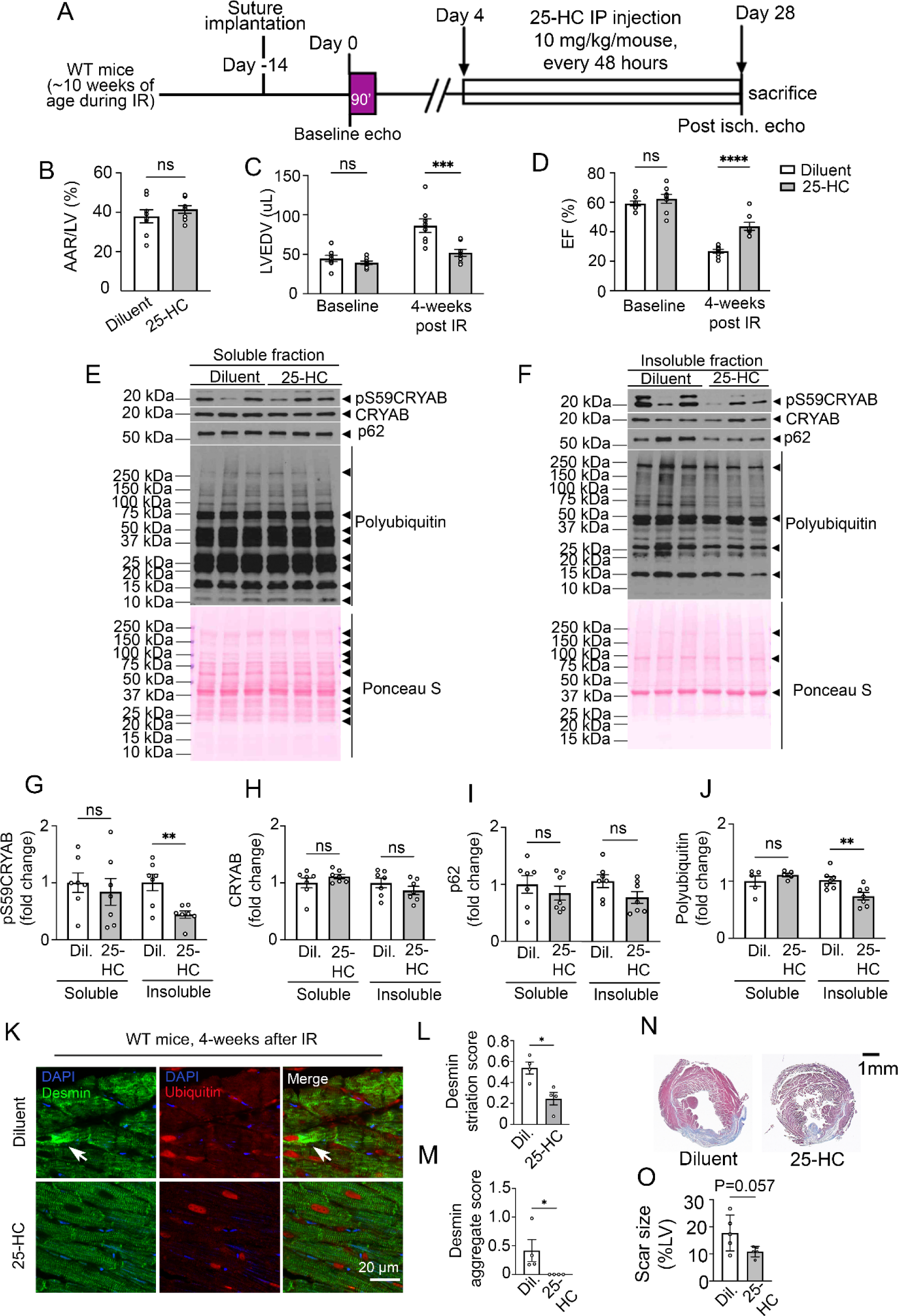
Treatment with 25-hydroxycholesterol rescues adverse LV remodeling after IR injury. **A.** Schematic depicting experimental strategy for closed-chest IR modeling (90 miutes of ischemia followed by reperfusion) in male wild-type mice followed by intraperitoneal administration of 25 hydroxycholesterol (10 mg/kg/mouse, every 48 hours) or diluent, initiated as day 4 after IR injury. **B-D)** Quantitative analyses of area-at-risk (AAR, B), left ventricular end-diastolic volume (LVEDV, C) and LV ejection fraction (EF (%), D) prior to and at 4 weeks after IR injury in mice treated as in A. *** denotes P< 0.001 and **** denotes P <0.0001 by t-test. ‘ns’ indicates not significant. **E, F)** Immunoblots depicting total CRYAB, pS59-CRYAB, p62 and polyubiquitinated protein expression in NP-40 soluble (E) and insoluble (F) fraction in myocardium of mice 4-weeks after IR injury and treated with diluent or 25-HC. **G-J)**. Quantitative analyses of pS59-CRYAB (G), total CRYAB (H), p62 (I) and polyubiquitinated proteins (J) in NP-40 soluble and insoluble fraction in myocardium of mice 4-weeks after IR injury and treated with diluent or 25-HC. **K**) Representative images with immunofluorescence staining for desmin and polyubiquitinated proteins in the remote myocardium from wild-type mice 4 weeks after closed-chest IR modeling followed by intraperitoneal administration of 25 hydroxycholesterol (10 mg/kg/mice in every 48 hours) or diluent. **L, M**) Quantitative evaluation of desmin localization with striation score (L) and aggregated desmin (L) in mice treated as in A. For striation scoring, normal localization of proteins got scored as 0; and abnormal striation or mis-localization of proteins was scored as of 1. For scoring aggregates, absence of aggregates was scored as 0 and presence of aggregates was scored as 2. * denotes P <0.05 by t-test. **N, O**) Masson’s trichrome stained left ventricular sections (N) demonstrating presence of scar at 4 weeks post-IR injury in mice treated as in A with quantitation of scar size (O). P value shown is by t-test.

## Discussion

Protein-aggregates are a hallmark of neurodegenerative diseases and are implicated in pathogenesis of rare genetic and amyloid cardiomyopathies. Our work demonstrates a mechanistic role for protein aggregation in the pathogenesis of ischemic cardiomyopathy, a prevalent condition which is the leading cause of heart failure worldwide. These findings implicate a broader pathogenic role for ‘condensatopathy’ as the underlying mechanism in cardiac disease beyond that observed in rare genetic cardiomyopathy such as the condition provoked by a mutant RBM20 protein (43). A mechanistic framework that emerges is that CRYAB, a highly enriched chaperone protein in cardiac myocytes undergoes phase separation into ‘liquid-like’ dynamic condensates under physiologic conditions (Figure 9). Sustained stress, as with development of ischemic cardiomyopathy, results in increased phosphorylation of CRYAB at serine-59 (Figure 1 and S2), which alters its phase separation behavior (Figures 2 and 3). Phospho-mimetic serine to aspartic acid change at position serine-59 in CRYAB induces ‘gel-like’ condensates with reduced dynamicity and increased interaction with clients such as desmin, triggering aggregate formation and cytotoxicity and sequestration of desmin in protein-aggregates (Figures 4-6). Mice harboring non-phosphorylatable alanine instead of serine at position 59 in CRYAB are partially protected from post-myocardial infarction left ventricular systolic dysfunction and desmin mis-localization with reduced scar size at 4 weeks post-MI (Figures 5 and 6). Also, reducing the overall abundance of pS59-CRYAB by 25-hydroxycholesterol treatment post-myocardial infarction, reduces protein aggregates with preserved desmin localization and attenuates ischemic cardiomyopathy (Figures S9, 7 and 8). These findings underscore the notion that targeting the mechanisms of altered phase separation, i.e. abnormal condensate behavior of CRYAB, is a clinically translatable strategy to ameliorate pathology in ischemic cardiomyopathy.

**Figure 9:**
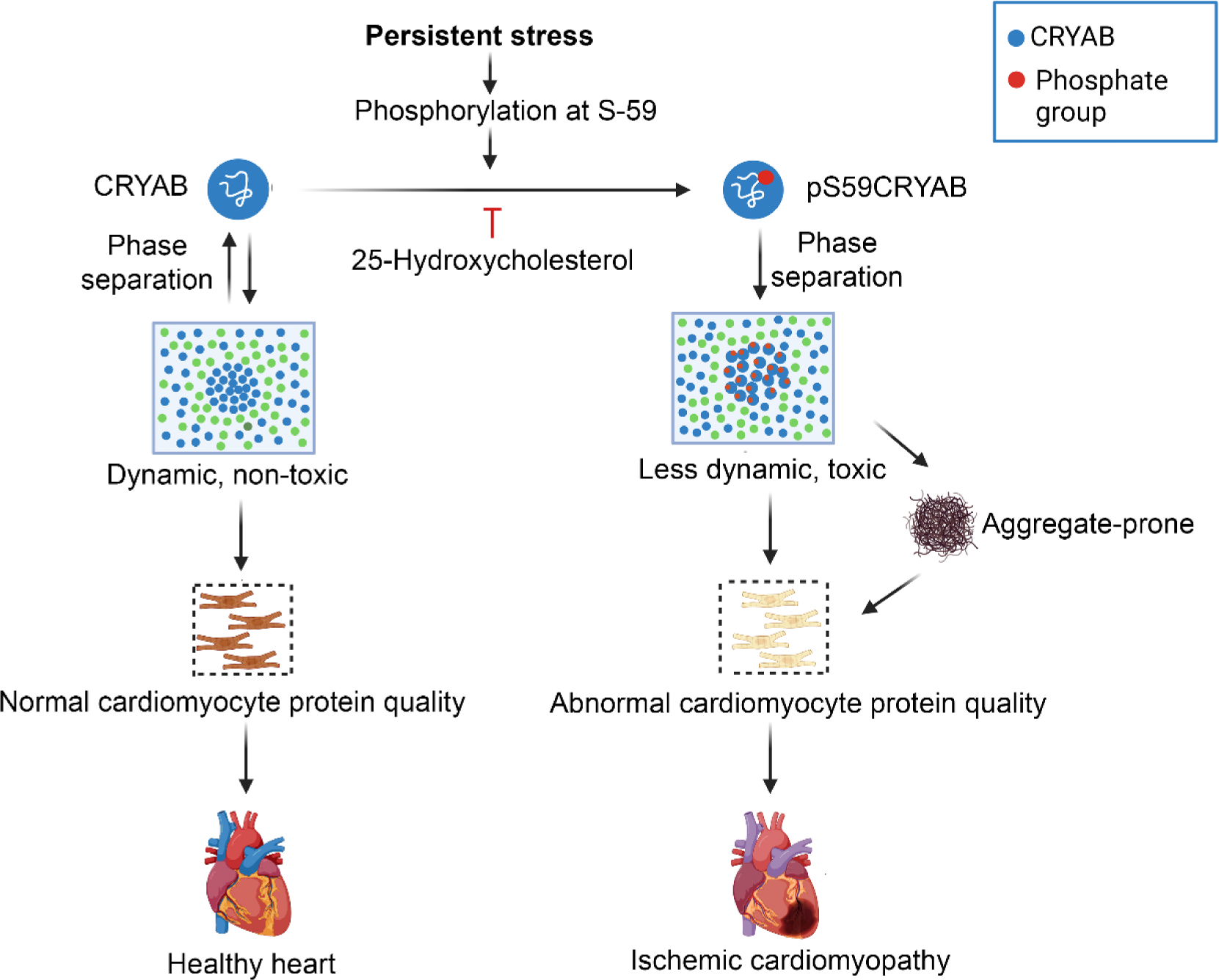
Schematic depicting consequences of serine-59 phosphorylation on phase separation of CRYAB in ischemic cardiomyopathy. CRYAB undergoes dynamic phase separation into condensates in physiology with maintenance of cardiac myocyte health in homeostasis. Persistent stress, as observed with myocardial ischemia reperfusion injury (or with human disease-causing R120G mutation in CRYAB protein, not shown), results in phosphorylation at serine-59. Phosphorylated CRYAB at serine-59 (pS59-CRYAB) undergoes phase separation into condensates with reduced dynamicity and increased toxicity. pS59-CRYAB partitions into the aggregate-rich NP-40 insoluble fraction with segregation of its client proteins, such as desmin, α-actinin and actin in aggregates within cardiac myocytes in ischemic cardiomyopathy. Inhibiting phosphorylation of CRYAB at serine-59 prevents it from becoming aggregate-prone and attenuates development of cardiomyopathy in the setting of IR injury. Image prepared with *BioRender* software.

Phase separation of proteins (along with other biomolecules) and their regulation by post-translational modifications has emerged as a critical mechanism underlying the control of spatiotemporal organization of cellular components in physiology (44, 45). Cellular structures, such as P-bodies and stress granules are formed by liquid-liquid phase separation and retain dynamic flux across the boundaries to facilitate rapid formation and reversibility (45). Interestingly, long-lasting subcellular structures, such as the nucleolus and centromeres that are less dynamic are also formed and maintained through phase separation and are conceived as more ‘gel-like’ (46–48). Dysregulation of the physiologic role of biomolecular condensates has been implicated in the pathogenesis of cancer, neurodegenerative and infectious diseases (28); but its role in human cardiac pathology has not been widely explored. A carefully performed experimental analysis of the R636S variant of human RNA-binding motif protein-20 (RBM20) that has been causally linked to development of dilated cardiomyopathy, revealed dysregulation of RNA-protein condensates in the cytosol as the pathogenic mechanism (43). Recent experimental studies identified that HIP-55 (hematopoietic progenitor kinase 1–interacting protein of 55 kDa), a negative regulator of β-adrenoceptor signaling, undergoes phase separation to form condensates and reduced phosphorylation dysregulates HIP-55 condensates drives its aggregation and cardiac dysfunction in mice challenged with isoproterenol (49). Our observation of CRYAB phosphorylation induced-condensatopathy post-myocardial infarction is the first to implicate a role for dysregulated condensates in a common etiology for heart failure.

Our data suggest that reversible stress-induced phosphorylation of CRYAB is likely a mechanism to regulate protein-protein interactions with client proteins by altering the dynamicity of the resultant condensates, and confers cytoprotection or cytotoxicity, depending upon the cellular context. Under acute stress, p38 kinase activation and resultant serine-59 phosphorylation of CRYAB is likely to be temporally limited and reversible with inactivation of p38 when stress abates, facilitating increased interaction with its client proteins such as desmin by reducing dynamicity of condensates, as a protective mechanism only while the stress lasts, akin to the physiology observed with stress granules (44). Indeed, CRYAB is rapidly phosphorylated at serine-59 with ischemia and localizes to the Z-discs in cardiac myocytes (31, 50) where it interacts with desmin with increased affinity under low pH (31); and binds tightly to its other client proteins including Titin, in a manner resistant to solubilization with urea (51). Also, CRYAB was noted to be transcriptionally upregulated in differentiating neural progenitor cells and localizes to aggregates to sequester mis-folded proteins; and siRNA-mediated knockdown of CRYAB prevents sequestration of mis-folded proteins with reduced cell viability (52). Analogously, overexpression of CRYAB with a phosphomimetic serine to aspartic acid change at position 59 conferred cytoprotection against hypoxic stress in isolated cardiac myocytes (41). In contrast, persistent stress with development of ischemic cardiomyopathy induces sustained serine-59 phosphorylation with persistent p38 kinase activation (53), whereby the pS59-CRYAB becomes a ‘toxic’ protein and requires sequestration within aggregates to mitigate toxicity. Indeed, prior studies have shown a high propensity for pS59-CRYAB to partition in a detergent-insoluble fraction in aggregates, termed Rosenthal bodies, in brains of patients with Alexander’s disease, and in Alzheimer’s disease (54), in the setting of persistent disease pathology.

The propensity of CRYAB to phase separate through intermolecular interactions to form higher order structures is likely to be facilitated by its intrinsically disordered regions, consistent with the reported heterogeneity of CRYAB multimers ranging from 10-40 subunits (55) indicating its polydisperse nature (56). Also, in rigorous biophysical studies, the N-terminal region of human CRYAB (residues 1-67) was found to exist in different states consistent with the notion of being intrinsically disordered and to contribute to the observed heterogeneity of the CRYAB protein multimers (57). These studies also predict a likely effect of S59 phosphorylation via interactions of the ‘β2’ region (Ser59-Thr63) with other regions of CRYAB, both intramolecularly as well as intermolecularly (57), on the composition of CRYAB multimers. Intriguingly, we have observed that 25-hydroxycholesterol inhibits phosphorylation of CRYAB at serine-59, as a novel mechanism to prevent aggregation of CRYAB. Our data also uncover this as a novel mechanism of action whereby 25-hydroxycholesterol drives increased solubility of the R120G protein with disruption of aggregates, complementary to its ability to bind CRYAB as suggested by molecular docking studies (40). Indeed, our studies with the phosphorylation-deficient mutant confirm that phosphorylation of CRYAB at serine-59 is essential for formation of aggregates with the R12OG mutant. It is also conceivable that 25-hydroxycholesterol acts as a hydrotrope to disrupt ‘gel-like’ condensates of R120G by preventing its multimerization, akin to a role for ATP (58).

Our study is the first to mechanistically implicate protein aggregation in development of ischemic cardiomyopathy. Prior work in myocardial samples from human ischemic cardiomyopathy has demonstrated increased abundance of short fibrils of desmin (∼190kD) which mimic pre-amyloid oligomers detectable with A11 staining in patients with dilated cardiomyopathy and mice with transgenic expression of R120G mutant CRYAB (16). These studies demonstrate presence of desmin in protein aggregates in mice subjected to pressure overload (16) or dogs modeled for rapid pacing-induced cardiomyopathy (59) and observe that desmin cleavage and phosphorylation are associated with formation of cardiac aggregates and pre-amyloid oligomers in the observed cardiomyopathy. Our work suggests a mechanism for desmin localization to aggregates by increased interaction with phosphorylated CRYAB which is aggregate-prone (Figure 5). It is also conceivable that post-translational modifications of CRYAB client proteins, such as desmin also alters their phase separation properties to drive protein aggregation and cause cardiac dysfunction, a premise that will require experimental validation.

Our discovery of the regulation of phase separation behavior of a cardiac myocyte-enriched chaperone protein, CRYAB, by a stress-induced post-translational modification, offers a therapeutic opportunity to explore and harness the beneficial signaling conferred by CRYAB and other chaperone proteins that are critical for cardiac myocyte protein quality control. These observations support a novel paradigm that dysregulation of biomolecular condensates, as observed in rare genetic cardiomyopathies triggered by the R636S mutation in RBM20 (a striated muscle-specific nuclear alternative splicing factor (43)), and with the R120G mutation in CRYAB (*vide supra*), may be pathogenic in common etiologies of cardiomyopathy; opening the door for targeting biomolecular condensates to develop sarcomere-targeted therapies for highly prevalent causes of heart failure.

## Methods

### Genetically modified mice

For generating mice with serine to alanine change (S59A) and serine to aspartic acid change (S59D), we employed CRISPR/Cas9 genome editing. A single active gRNA was used to introduce point mutations in the CRYAB locus of B6/CBA hybrids using homology-directed repairs (HDR) after the pro-nuclear introduction of both gRNA and Cas9 was done by electroporation. ZE electroporation was performed in four sessions where 2 different mice colonies, S59A (AGC to GCC) and S59D (AGC to GAC), were generated with a single electroporation mix (i.e. both ssODNs in the same mix). The presence of the CRYAB point mutations was confirmed in each generation of mice colony by deep sequencing. We used both male and female mice from the seventh generation of backcrosses into the C57BL/6J strain (JAX strain 000664) for the experiments. Mice of both sexes were studied, unless otherwise specified. No significant differences were observed between sexes for the primary phenotype, whereby the data for both sexes were combined for presentation. All observers were blinded.

### Human Heart tissues

De-identified frozen heart tissue samples were obtained from the Human Heart Tissue Bank at the University of Pennsylvania (as described in Supplemental methods), and formalin-fixed paraffin-embedded left ventricular myocardial tissue was obtained from the Translational Cardiovascular Biobank and Repository at Washington University School of Medicine. The hearts were procured from two separate patient groups: non-failing brain-dead organ donors with no history of heart failure (Donor, whose hearts were screened but not selected for transplantation) and heart transplant recipients with advanced ischemic cardiomyopathy (ICM), that were obtained at the time of orthotopic heart transplantation.

Clinical characteristics of the subjects providing human tissue for these studies are summarized in Supplemental Table S1.

### Closed-chest cardiac ischemia-reperfusion modeling

Mice were subjected to reversible left anterior descending coronary artery ligation for 90 min followed by reperfusion, in a closed-chest procedure, as described (60). Overall surgical mortality was <5%. All surgeries were performed by two surgeons (C.J.W. and J.N.). In a cohort of animals, cardioplegic solution was injected retrogradely through the aorta *in-situ* at 24 hours post-induction of ischemia, followed by injection of Evans Blue dye, and sectioning of the left ventricle in slices, which were incubated in triphenyl tetrazolium chloride (TTC) for 30 min at 37°C. For infarct size calculation, TTC-stained slices were imaged and infarct area quantified as non-TTC-stained area as a ratio of total non-blue myocardial area, as previously described (61).

### Echocardiography

2D–directed M-mode echocardiography was performed using a Vevo 2100 Imaging System (VisualSonics, Toronto, Canada) equipped with a 30-MHz linear-array transducer, as we have previously described (60, 62). Mice were anesthetized with avertin for studies obtained in unstressed mice and before I/R modeling (pre-ischemia echo study). Echocardiography during ischemia was performed under isoflurane anesthesia and at four weeks post ischemia-reperfusion injury was performed under avertin anesthesia. Cardiac images were obtained by a handheld technique. Area-at-risk, left ventricular dimensions, wall thickness, heart rate, fractional shortening, EF and volume calculations were performed by a blinded echocardiographer as described (60). Histologic assessment with hematoxylin and eosin staining, and assessment of myocardial fibrosis with Masson’s trichrome staining and transmission electron microscopy, was performed as previously described (18).

### Neonatal cardiac myocyte isolation

Neonatal rat cardiac myocyte isolation was performed using the technique we have described with the Worthington Neonatal Cardiomyocyte Isolation System (CAT# LK003300) (61). Hearts were harvested from one-day old neonatal rats and were subjected to trypsin digestion in a final concentration of 50 μg/ml in HBSS for 16-18 hours at 4°C after removal of the atria. Collagenase digestion (type II collagenase; 300 U/ml; Worthington) was conducted at 37°C for 45 min. Cardiomyocytes were seeded on collagen-coated four well chamber slides (Laboratory Tek) at a density of 10^5^ cells per cm^2^. On the 2nd day, the culture medium was changed to the Rat Cardiomyocyte Culture Medium (Cell applications INC, CAT#R313-500) prior to immunofluorescence staining.

### Adenoviral studies

Adenoviral vector encoding for CRYAB-R120G (18) was employed, as previously described.

### Biochemical fractionation into soluble and insoluble fractions

We subjected cardiac tissues obtained from human hearts and from the remote left ventricular myocardium of mice subjected to ischemia-reperfusion injury (non-injured basal one-third); and cell extracts from neonatal rat cardiac myocytes to obtain soluble-insoluble fractions as previously described (18). Heart tissue was mechanically homogenized in homogenization buffer (0.3 M KCl, 0.1 M KH_2_PO_4_, 50 mM K_2_HPO_4_, 10 mM EDTA, 4 mM Na Orthovanadate, 100 mM NaF, 1X Protease inhibitor, pH to 6.5). Homogenized samples were passed through mesh basket on ice, followed by collection of the lysate run-through which was incubated on ice for 30 minutes. An aliquot was transferred to another Eppendorf tube and 10% NP-40 was added to for a final concentration of 1% NP-40. Samples were then incubated on ice for 30 minutes and spun at 16,000g for 15 minutes at 4°C. Supernatant was collected as soluble fraction. The pellet was washed 3 times with cold PBS (following addition of 1ml PBS to each pellet, and spin down at 16,000g for 10 minutes) followed by resuspension in 1% SDS, 10mM Tris buffer to generate the insoluble fraction.

### Co-Immunoprecipitation

Frozen hearts were placed in RIPA buffer (Cell Signaling 9806) containing Protease and Phosphatase Inhibitor (Thermo Fisher 78442) and homogenized with a mechanical homogenizer. Tissue lysis was pelleted at 600g for 5min. (Both supernatant and pellet were used for Co-IP). The pellet was dissolved with PBS containing 1% SDS and 5mM EDTA, and sonicated. The solution was centrifuged at 16000g for 5 min and the supernatant were diluted by 10 with PBS containing 5mM EDTA. Protein A/G Sepharose (Abcam ab193262) were incubated with 5μg antibody or IgG in PBS for 30 min at room temperature with rotation. The immunocomplex was added to the lysis solution after 15min incubation with rotation at room temperature. The beads were washed with PBS for four times. The proteins were eluted by boiling in SDS sample buffer and analyzed by western blot. The antibodies used in this experiment were: Anti-desmin (Desmin (D93F5) XP^®^ Rabbit mAb #5332), anti-CRYAB (Abcam, CAT# ab13496).

### Immunofluorescence analysis

We performed immuno-histochemistry on cells and myocardial tissues as we have previously described (18). Primary cultures of neonatal rat cardiac myocytes were fixed in 100% cold methanol for 20 minutes, followed by blocking with 1% normal serum in PBS for 1 hour at room temperature. Primary antibodies used were as follows: anti-desmin (Santa Cruz Biotechnology, Inc, CAT#SC-7559), anti-ubiquitin (Sigma-Aldrich, CAT#04-262), anti-actin (Millipore Sigma, CAT#A2066), anti-α-actinin (Abcam, CAT#ab9465), anti–αB-crystallin polyclonal antibody (ENZO Life, ADI-SPA-223-F) with overnight incubation at 4°C. Paraffin-embedded heart sections (4 µm thick) were subjected to de-paraffinization using Xylene; serial 100%, 90%, 70% and 50% EtOH treatment followed by hydration using DI water and heat-induced epitope retrieval in Diva decloaker solution (Biocare medical, REF# DV2004MX). This was followed by blocking using 1% BSA (Sigma-Aldrich, CAT#A9647-100G), 0.1% Tween-20 (Sigma-Aldrich, CAT# P2287-500ML) in PBS (Corning, CAT# 21-040-CM) and 5% donkey serum. The slides were incubated overnight with primary antibody. Next day, after serial washes, samples were stained with secondary antibody and mounted with fluorescent 4’,6-diamidino-2- phenylindole mounting medium (Vector Labs, H-1200). For A11-staining (ThermoFisher, CAT#AHB005), the protocol was revised according to vendors instructions. The antigen retrieval solution was 0.1M glycine/PBS, at pH 3.5. Anti-oligomer A-11 antibody was used at 1-5 ug/mL concentration in 1:1mixture of blocking solution (1%BSA, 0.1% Tween-20 in PBS) and PBS. Confocal imaging was performed on a Zeiss confocal LSM-700 laser scanning confocal microscope using 40×/1.3 oil immersion objectives, and images were processed using the Zen black software.

### Striation and aggregate scoring

In order to quantitate the mis-localization and aggregation of CRYAB client proteins, namely desmin, actin and α-actinin we utilized the following scoring syste. 1) For striation scoring, normal localization of proteins got scored as 0; and abnormal striation or mis-localization of proteins was scored as of 1. 2) For scoring aggregates, absence of aggregates was scored as 0 and presence of aggregates was scored as 2. Then, all the individual scores of the cardiomyocytes in the total field of view were added and divided by total number of cardiomyocytes. We quantified at least 3 images per sample. Image acquisition and quantitation was done by different operators where scoring was done blindly.

### Immunoblotting

Immunoblotting was performed as previously described (18) with antibodies listed below. Specific antibodies employed are as follows: anti-SQSTM1/p62 antibody (Abcam, ab56416); αB-crystallin (CRYAB) polyclonal antibody (Enzo life sciences, CAT# ADI-SPA-223- F); Anti-αB crystallin (pS59) antibody (Abcam, CAT#ab5577) (63); Anti-αB crystallin (pS45) antibody (Abcam, CAT#ab5598) (63); Anti-ubiquitinylated proteins antibody, clone FK1 (Sigma-Aldrich, CAT#04-262) ; Anti-GAPDH antibody (Abcam, CAT#ab22555); Anti-Actin antibody (Sigma, CAT#A2066); anti-GFP antibody (Abcam, CAT#ab290), anti-Desmin antibody (Santa Cruz Biotechnology, CAT#SC-7559).

### Generation of crystallin constructs

pHR-mCh-Cry2WT (Addgene #10221) and pHR-FUSN-mCh-Cry2WT (Addgene #10223) as generated by Dr. Clifford Brangwynne (39), were obtained and verified by sequencing. For phase separation assay in the live cells, we have generated pHR- CRYAB-mCh-Cry2 construct using in-fusion® snap assembly kit (Takara, CAT#638945). CRYAB S59A, S59D and R120G+S59A mutations were introduced in pHR-CRYAB-mCh-Cry2 construct using QuikChange II site-directed mutagenesis kit (Agilent, CAT#200523). These constructs were subsequently cloned into pcDNA 3.1 mammalian vector for the expression in HEK293A cells. Constructs coding for eGFP-CRYAB, eGFP-CRYAB S59A, eGFP-CRYAB S59D, eGFP-CRYAB-R120G+S59A and eGFP-CRYAB-R120G+S19A+S45A+S59A were generated as described above in a pcDNA3.1 backbone for protein aggregation studies and cell death assessment.

### Assessment of protein aggregation and cell death

HEK293 cells were transfected with various GFP-tagged CRYAB constructs using Lipofectamine 3000 reagent (ThermoFisher, CAT#L3000) and cells imaged for GFP at 24 hours after transfection under Zeiss-700 confocal microscope for presence of protein aggregation. Cells were harvested at 48 hours for cell death assay. Cell death was assessed using a fluorometric assay with LIVE/DEAD™ Viability/Cytotoxicity Kit for mammalian cells (ThermoFisher, CAT#L3224), as we have previously described (18).

### Studies with optoDroplet constructs

CRYAB constructs were transfected into HEK 293A cells using Effectene transfection reagent (Qiagen, CAT#301425) and live-cell imaging was performed using 35-mm glass-bottom dishes at 24-hours after transfection using 40x oil immersion objective of the Nikon A1R confocal imaging system (equipped with 37°C stage) at the Center for Cellular Imaging (WUCCI) at Washington University in St. Louis School of Medicine. Cells were imaged with two laser wavelengths (488 nm for Cry2 activation and 561 nm for mCherry imaging) with laser power of 10% for the 488 nm. Various parameters were quantified using imageJ (NIH) by adjusting the threshold to ensure uniform cut-off values at the t=0 and t=300s time-point images. Average condensate area was determined by total area of the fluorescence signal divided by number of condensates before or after 300s of the blue light activation. Average number of condensates per cell was obtained by dividing total number of condensates divided by number of cells before or after 300s of the blue light activation.

### Studies with fluorescence recovery after photobleaching (FRAP)

FRAP was performed on OptoIDR constructs as described above while imaging with100x objective lens of the Nikon A1R confocal imaging system, using 488nm laser at 50% intensity and 561 nm laser at 50% intensity for 1 minute selecting a region of interest of ∼ 1 µm in diameter; and fluorescence recovery was monitored. Intensity traces were collected using imageJ and normalized to pre-bleaching intensity (set at 100%). Only condensates with reduction in intensity to <10% (as compared to pre-bleach) post-photobleaching were selected for further analysis.

### Studies with 25-hydroxycholesterol

25-hydroxycholesterol (CAT#H1015-100MG, Sigma-Aldrich) was dissolved in 100% ethanol (final concentration 20mg/mL) to make a stock solution. For in-vivo experiments, the stock solution was diluted in sterile PBS for injection. A dose of 10mg/kg or equivalent vehicle control was administered every other day via intraperitoneal injection beginning at day 4 after closed-chest ischemia-reperfusion injury for 3.5 weeks. Mice were weighed prior to injection and food intake was monitored through the experiment.

### Statistical analyses

Data presented as mean±SEM. All measurements were obtained from separate biological replicates. Statistical analyses were performed in GraphPad Prism version 9. Data were tested for assumptions of normality with Shapiro-Wilk normality test. Statistical significance was assessed with unpaired two-tailed Student’s t-test for comparison between two groups, or one-way or two-way ANOVA for assessing differences across multiple groups followed by post-hoc testing. A non-parametric test was performed if data were not normally distributed. A two-tailed P value of <0.05 was considered statistically significant.

### Study approval

All animal studies were approved by the IACUC at Washington University School of Medicine. Studies on human tissue were performed under an exemption by the IRB at Washington University School of Medicine as only de-identified human samples were used and these studies were deemed exempt by the IRB at Washington University School of Medicine.

## Author contributions

M.I., D.R.R., X.M., W.N., C.Z., L.F., J.T.M., C.J.W., H. N., Aa. D., R.C. and M.K., performed experiments, acquired, and analyzed the data; A.K., J.N. acquired and analyzed the data; B.R. provided critical reagents, interpreted the data and revised the manuscript; M.E.Y., M.A., S.S., K.B.M., D.F.C. and A.J., interpreted the data and revised the manuscript; K.M. and A.D. conceived the experiments, analyzed the data and drafted the manuscript. K.M. and A.D. supervised the work.

## Acknowledgements

The authors wish to thank Drs. Douglas L. Mann, Justin Hartupee and Kory Lavine from Washington University School of Medicine for providing fixed human myocardial tissue sections.

