## Supplementary Data for "Phosphorylation of CRYAB Induces a Condensatopathy to Worsen Post-Myocardial Infarction Left Ventricular Remodeling"

**Table of Contents for Supplementary Data**

**Supplementary Fig. S1: Desmin localizes to protein-aggregates and pre-amyloid oligomers are observed in human ischemic cardiomyopathy……………………………………. ** Page 2

**Supplementary Fig. S2: CRYAB phosphorylated at serine-59 partitions to the insoluble fraction in the myocardium of young adult C57BL6J WT mouse subjected to closed chest ischemia-reperfusion (IR) with development of ischemic cardiomyopathy…………**Pages 3, 4

**Supplementary Fig. S6: Serine to alanine change at position 59 prevent phosphorylation at this residue in CRYAB………………………………………………………………..** Page 8

**Supplementary Fig. S7: Crispr-Cas9 knock-in of phosphorylation-deficient serine to alanine change at position 59 (S59A) and phospho-mimetic serine to aspartic acid change (S59D) does not alter total CRYAB and desmin abundance………………………………..** Page 9

**Supplementary Fig. S8: 25-HC does not affect phase separation or dynamicity of CRYAB S59D…………………………………………………………………………………….** Page 10

**Supplementary Fig. S9: 25-HC treatment reduces overall pS59CRYAB in the WT C57BL6J mice subjected to IR injury………………………………………………………………** Page 11

**Table S1: Characteristics of individuals whose human heart samples were included in the study……………………………………………………………………………………..** Page 12

**Table S2: Morphometric and M-mode echocardiographic data for CRYAB WT, S59A homozygous knock-in, and S59D homozygous knock-in mice at 10 weeks of age…..** Page 13

**Supplementary Videos S1-S14…………………** Available as Separate files

Supplementary Figure S1

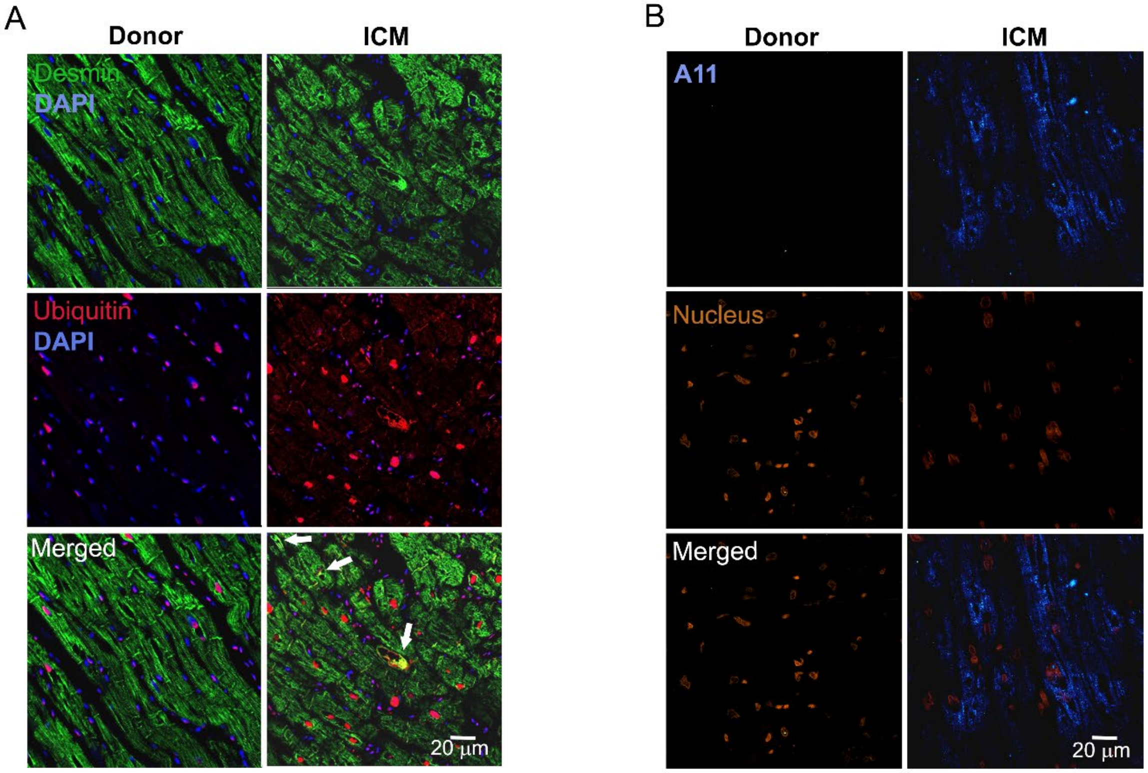

**Supplementary Fig. S1: Desmin localizes to protein-aggregates and pre-amyloid oligomers are observed in human ischemic cardiomyopathy. A.** Representative immunohistochemical images from left ventricular myocardium of individuals evaluated as controls (donor) or patients with end-stage ischemic cardiomyopathy (ICM) stained for desmin and polyUb. Arrows point to desmin, which is mis-localized from its physiologic location on Z-discs and intercalated discs in donor myocardium to protein-aggregates stained with polyUb antibody in ICM myocardium. DAPI stains nuclei. Representative of n=3 hearts/group. **B.** Immunohistochemical staining for anti-oligomer A11 antibody in donor and ICM heart samples, demonstrating presence of pre-amyloid oligomers structures in ICM samples (pseudo-colored blue, arrows). DAPI stained nuclei are pseudo-colored orange. Representative of n=5 hearts/group.

#### Supplementary Figure S2

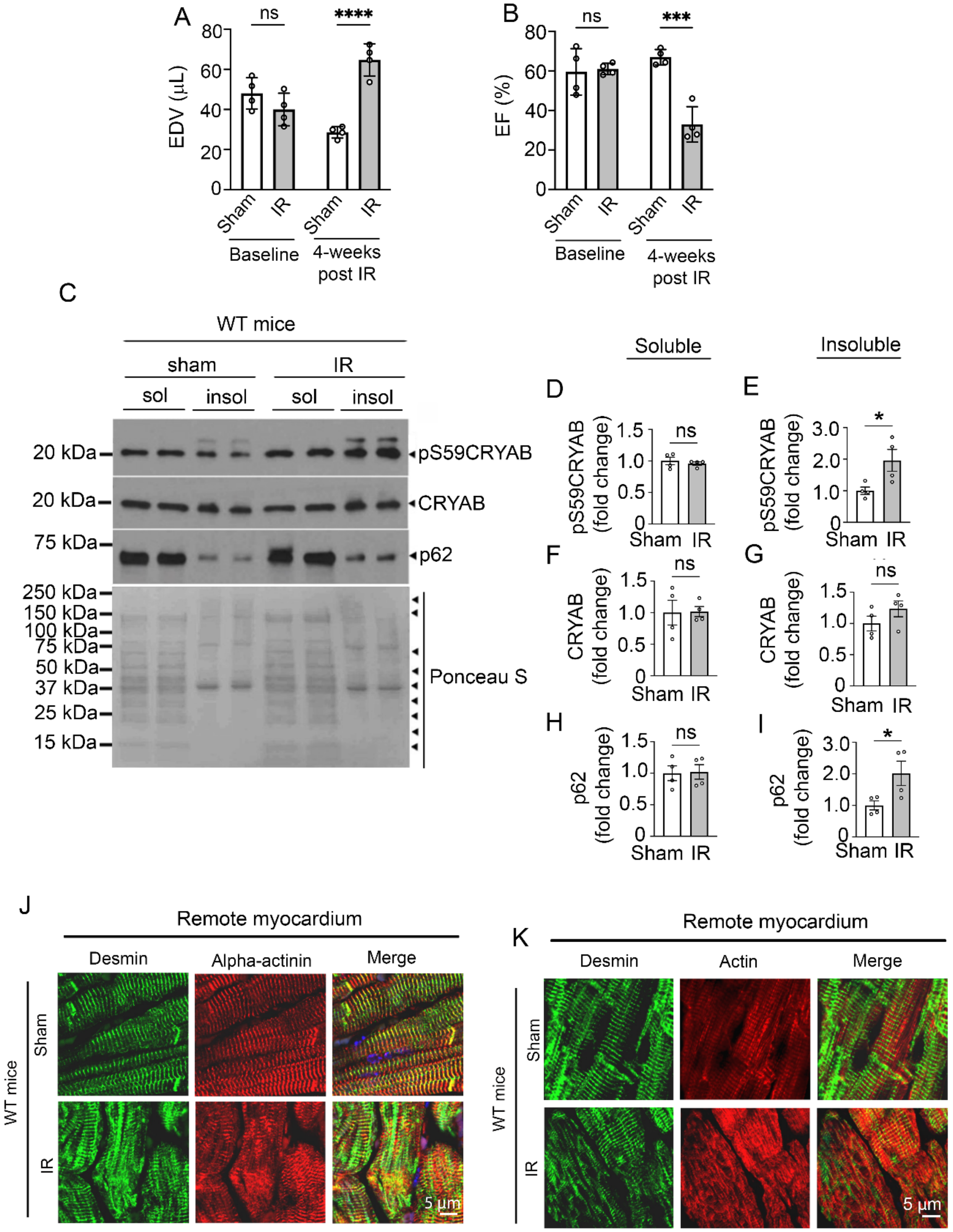

**Supplementary Fig. S2: CRYAB phosphorylated at serine-59 partitions to the insoluble fraction in the myocardium of young adult C57BL6J WT mouse subjected to closed chest ischemia-reperfusion (IR) with development of ischemic cardiomyopathy. A, B)** Left ventricular end-diastolic volume (EDV) (A) and left ventricular ejection fraction (EF, %) (B) at baseline and at 4-weeks after closed chest IR (90 minutes of ischemia followed by reperfusion) in male C57BL6J WT young adult mice that were subjected to IR injury or sham procedure. *** indicates P value < 0.001 and **** indicates P value <0.0001 by Tukey's post-hoc test after one-way ANOVA. **C-I)** Representative (C) immunoblot and quantitation depicting the abundance of pS59CRYAB (D, E), CRYAB (F, G) and p62 (H, I) in the soluble (*left* panels) and insoluble (*right* panels) biochemical fractions from the remote myocardium of the C57BL6J WT young adult mice collected 4 weeks after IR or sham procedure as in A, B. Ponceau S is shown as loading control. * indicates P<0.05 by t-test. **J, K)** Representative images demonstrating expression of desmin and α-actinin (J), and of desmin and actin (L) in the myocardium from C57BL6J mice 4 weeks after being subjected to IR injury. DAPI stains nuclei.

Supplementary Figure S3

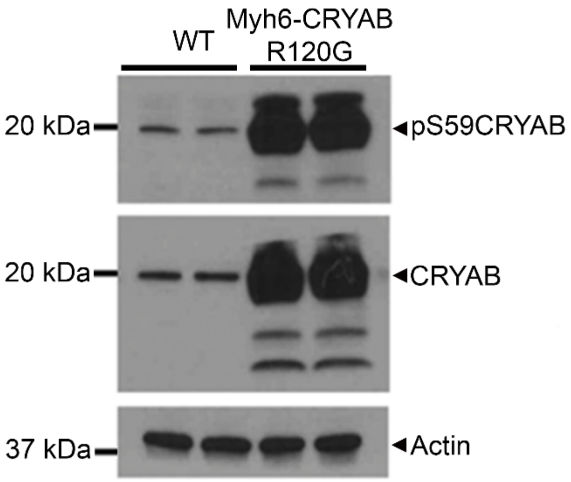

**Supplementary Fig. S3: R120G protein is prominently phosphorylated at S59**. **A)** Representative image showing expression of CRYAB and pS59CRYAB in crude extracts from 40 week-old Myh6-CRYABR120G mouse hearts or from C57BL6J WT mouse hearts as control. Actin is shown as the loading control.

#### Supplemental figure S4

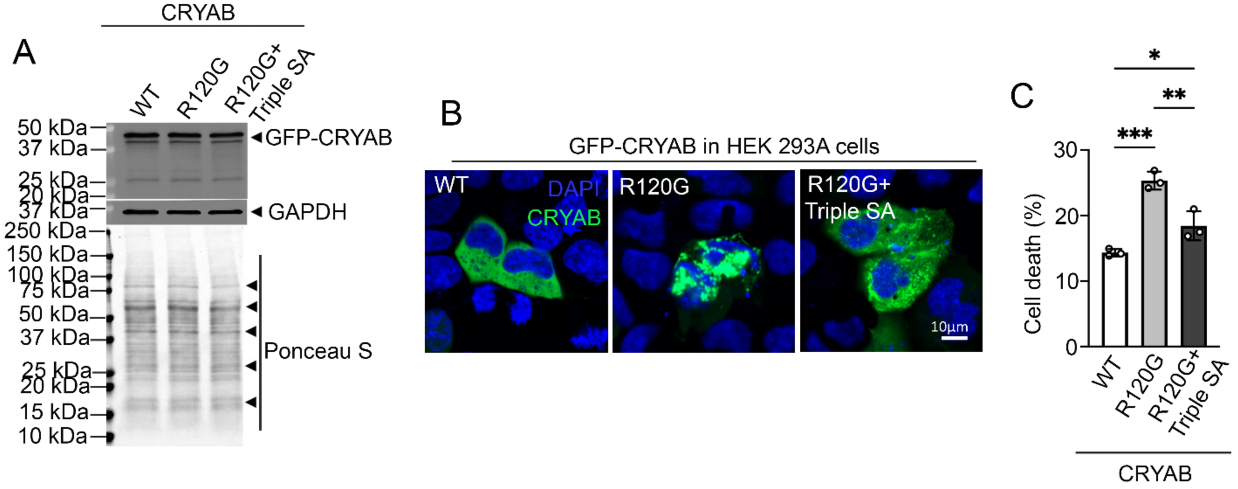

**Supplementary Fig. S4: Preventing phosphorylation at all three serine residues in CRYAB (S19, S45 and S59) reduces cell death in CRYAB R120G. A**) Immunoblot (A) demonstrating expression of GFP-fusion proteins in HEK293A cells transfected with GFP-tagged wild-type CRYAB, R120G mutant or the R120G and S19, S45, S59A triple mutant proteins. **B, C)** Representative immunofluorescence images (B) for detection of protein-aggregates with quantitation (C) of % cell death. * Indicates $p<0.05$, ** indicates $p<0.01$, and *** indicates $P<0.001$ by Tukey's post-hoc test after one-way ANOVA. Nuclei are blue (DAPI).

#### Supplementary Figure S5

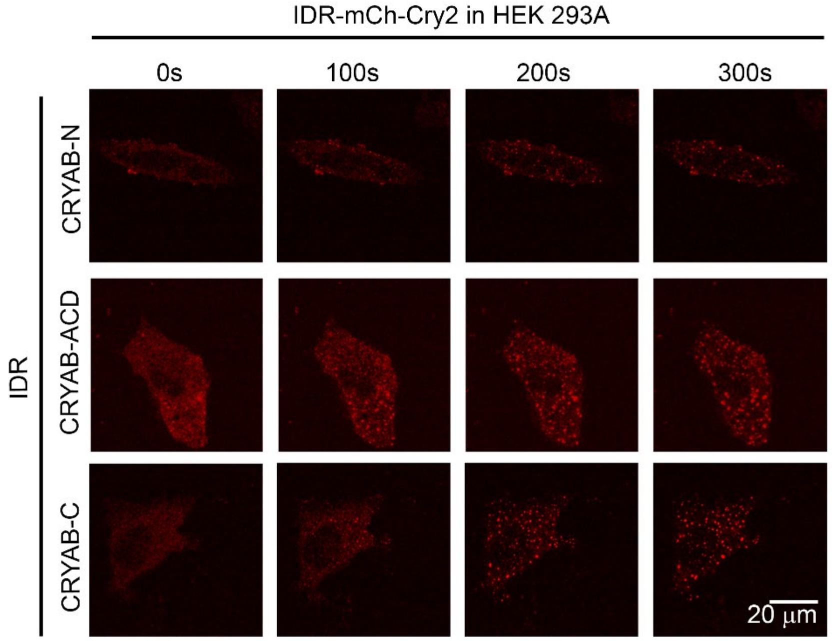

**Supplementary Fig. S5: N-terminus, ACD, and C-terminus domains of CRYAB undergo phase separation.** Representative time lapse images at t=0s, 100s, 200s, and 300s after light activation in HEK293A cells transfected with constructs generated with CRYAB N-terminus, Alpha-crystallin domain (ACD), and C-terminus domains as the 'IDR' in the optoIDR constructs.

Supplementary Figure S6

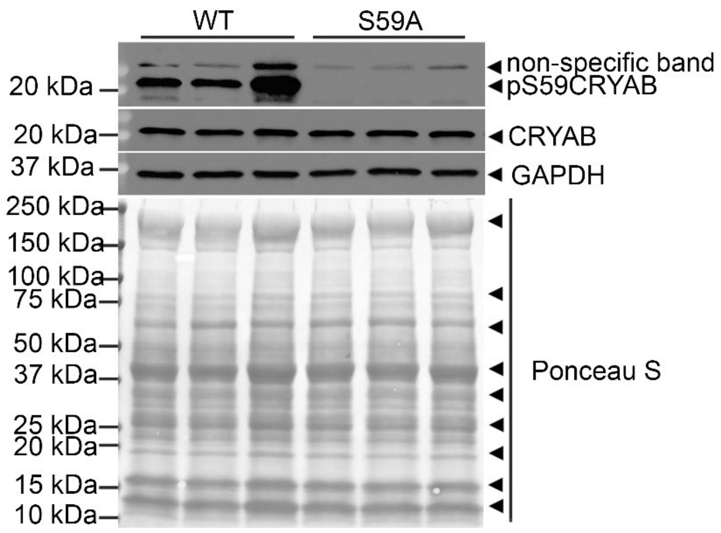

**Supplementary Fig. S6: Serine to alanine change at position 59 prevents phosphorylation at this residue in CRYAB.** Representative immunoblot demonstrating loss of immunodetectable band with an antibody that recognizes CRYAB phosphorylated on serine 59 (pS59-CRYAB), total CRYAB and GAPDH in myocardial extracts from mice homozygous for alleles bearing knock-in of alanine residue at this position. The specific band indicating pS59-CRYAB is indicated. Ponceau S is shown as loading control.

#### Supplementary Figure S7

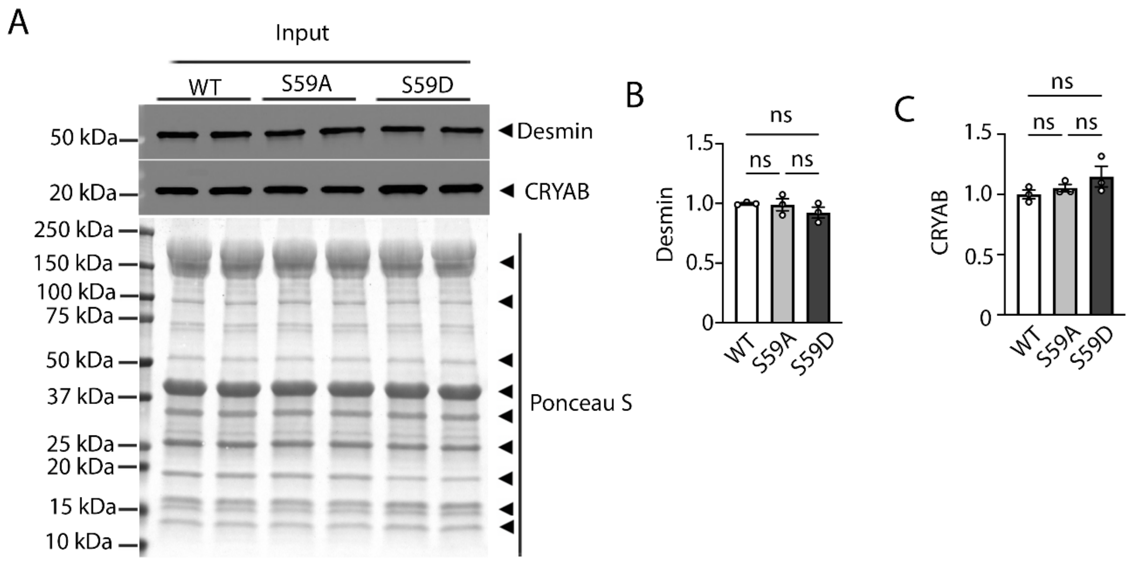

**Supplementary Fig. S7: Crispr-Cas9 knock-in of phosphorylation-deficient serine to alanine change at position 59 (S59A) and phospho-mimetic serine to aspartic acid change (S59D) does not alter total CRYAB and desmin abundance. A-C)** Representative immunoblot demonstrating expression of CRYAB and desmin in myocardial extracts from young adult mice homozygous for S59A or S59D CRYAB alleles or bearing wild-type CRYAB (WT) with quantitation of desmin (B) and CRYAB abundance (C) both expressed as fold over WT. 'ns' indicates no statistically significant differences were noted by one-way ANOVA analyses. Ponceau S is shown as loading control.

### Supplemental figure S8

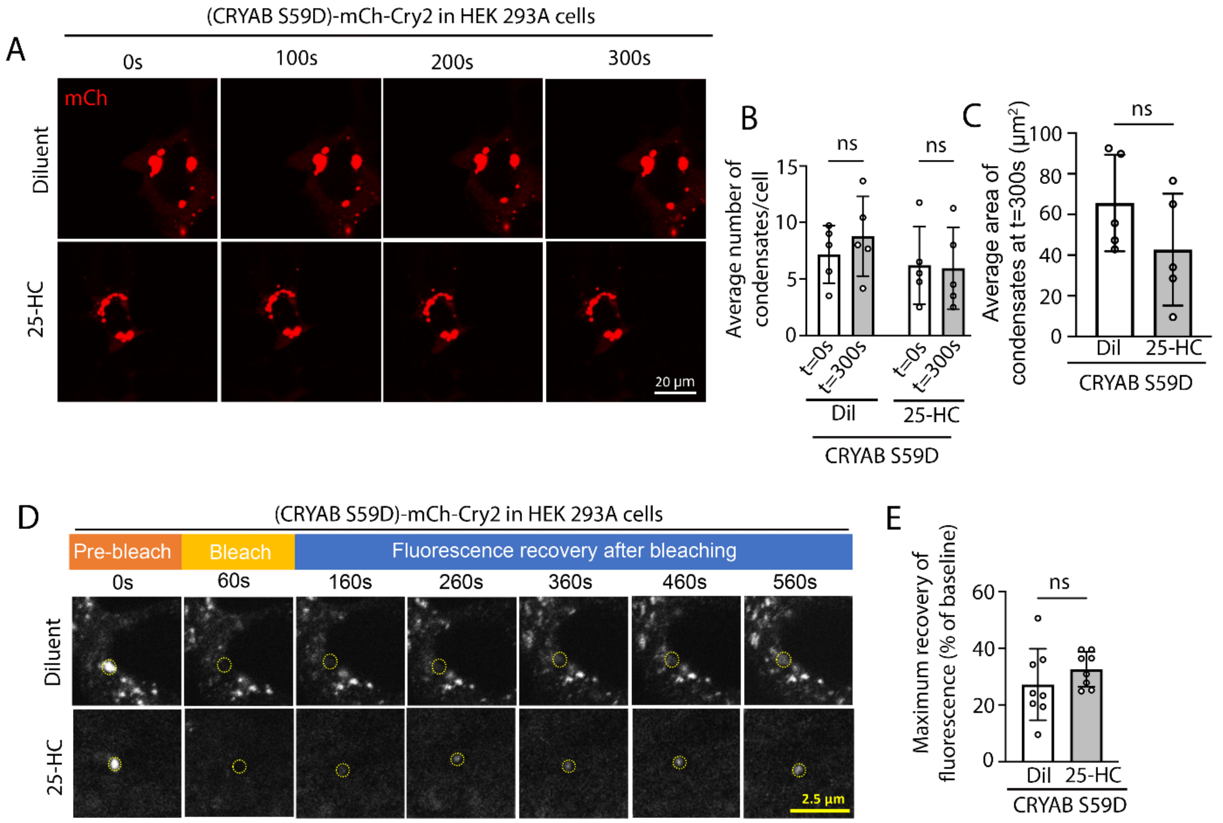

**Supplementary Fig. S8: 25-HC does not affect phase separation or dynamicity of CRYAB S59D. A)** Representative time-lapse images at t=0s, 100s, 200s, and 300s after light activation in HEK293A cells transfected with constructs generated with CRYAB S59D, the phospho-mimetic mutant as the 'IDR' in the optoIDR constructs. **B**) Average number of condensates/cell at t=0 vs. t=300s in cells treated as in E. 'ns' indicates not significant by t-test. **C)** Average area of condensates/cell at t=300s in cells treated in A. 'ns' indicates not significant by t-test. **D)** Representative images demonstrating recovery of fluorescence after photobleaching in HEK 293A cells transfected with mCherry-Cry2 fused optoIDR constructs generated with CRYAB S59D, the phospho-mimetic mutant. Representative images demonstrate area of photobleaching (marked with a dotted circle) prior to (pre-bleach), immediately after, and at 100, 200, 300, 400 and 500 seconds (s) after photobleaching was terminated. Intensity at various time points is depicted as a fraction of intensity prior to bleaching (set at 100%). **E)** Quantitation of fluorescence recovery (maximum minus immediately post-bleach) in CRYAB S59D variants indicated in D. 'ns' indicates not significant by t-test.

Supplemental figure S9

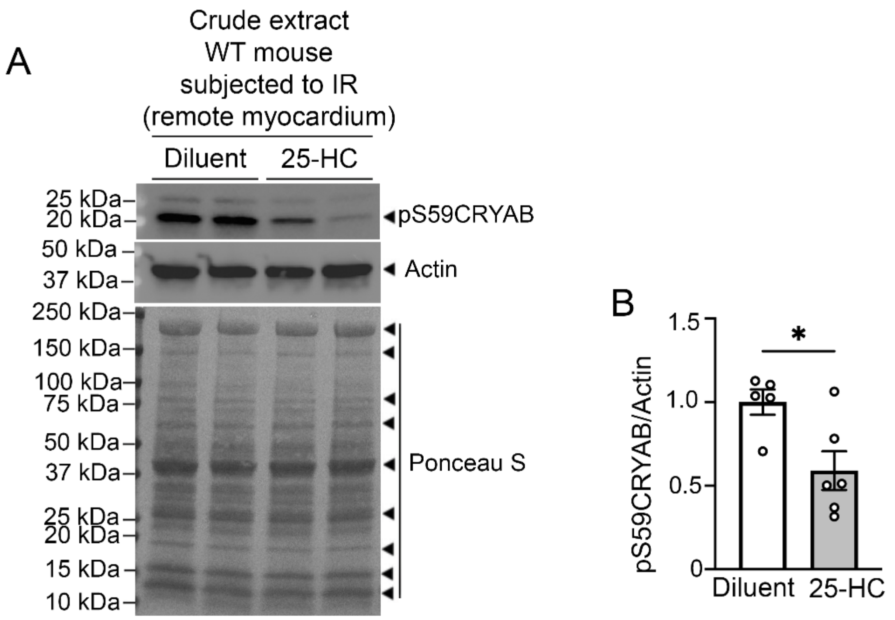

**Supplementary Fig. S9: 25-HC treatment reduces overall pS59CRYAB in the WT C57BL6J mice subjected to IR injury. A)** Representative immunoblot (A) and quantitation (B) of pS59CRYAB in crude extracts of remote myocardium from WT C57BL6J young adult male mice that were treated with 25-HC or diluent after IR injury as shown in Figure 8A. * indicates $p<0.05$ by t-test. Actin and Ponceau S were used as loading controls.

**Supplementary Videos S1-S14.** Representative videos depicting light induced phase separation of indicated optoIDR constructs, along with positive control (FUS-N) and negative control (Cry2).

**Supplementary tables**:

**Table S1: Characteristics of individuals whose human heart samples were included in the study.**

| **Individual Characteristics** | **Ischemic Cardiomyopathy (n=8)** | **Non-failing Donor (n=8)** | **Statistical Comparison** |
|---|---|---|---|
| Male Gender/Total Sample (N) | 5/8 | 6/8 | NS |
| Age (years) | 57 ± 3 | 52 ± 2 | NS |
| BMI | 27.79 ± 1.02 | 29.27± 3.74 | NS |
| LV Mass Index (g/kg) | 175.97 ± 3.76 | 105.38 ± 1.86 | P<0.0001 |
| LVEDD (cm) | 6.32 ± 0.28 | 4.03 ± 0.36 | P<.01 |
| LVEF (%) | 22.53 ± 3.85 | 60.13 ± 2.61 | P<0.0001 |
| History of Diabetes | 2/8 | 2/8 | NS |
| History of HTN | 4/8 | 1/8 | NS |
| History of ACE-I/ARB use | 4/8 | 0/8 | NS |
| History of Beta-blocker use | 4/8 | 0/8 | NS |

All data for continuous variables are mean±SEM. P values reported are by two-tailed t-test for continuous variables and by Fisher's exact test for categorical variables. NS indicates not significant.

**Table S2: Morphometric and M-mode echocardiographic data for CRYAB WT, S59A homozygous knock-in, and S59D homozygous knock-in mice at 10 weeks of age.**

| | CRYAB WT (n=6) | CRYAB S59A (n=8) | CRYAB S59D (n=8) |
|---|---|---|---|
| **BW (g)** | 22.95 ± 1.61 | 22.28 ± 1.06 | 21.09 ± 1.39 |
| **HW/BW (mg/g)** | 4.53 ± 0.11 | 4.71 ± 0.16 | 4.78 ± 0.07 |
| **HW/TL (mg/mm)** | 6.50 ± 0.33 | 6.29 ± 0.18 | 6.36 ± 0.39 |
| **LVIDd (mm)** | 3.29 ± 0.09 | 3.13 ± 0.06 | 3.20 ± 0.10 |
| **LVIDs (mm)** | 1.76 ± 0.10 | 1.62 ± 0.04 | 1.75 ± 0.09 |
| **FS (%)** | 46.65 ± 1.81 | 48.43 ± 0.79 | 45.69 ± 1.19 |
| **LV mass (mg)** | 63.41 ± 2.98 | 57.37 ± 1.55 | 63.34 ± 4.41 |
| **HR (beats per min)** | 625± 14 | 593 ± 8 | 608 ± 11 |

Data represent mean ± SEM. No significant differences were found between groups in ordinary one-way ANOVA. BW= Body Weight, HW= Heart Weight, TL= Tibia Length, LVIDd= Left Ventricular Internal Diameter End Diastole, LVIDs= Left Ventricular Internal Diameter End Systole, FS= Fractional Shortening, LV mass= Left-Ventricular mass based on M-mode echocardiography measurements, HR= Heart rate.
